# Continental-scale relationships of fine root and soil carbon stocks hold in grasslands but not forests

**DOI:** 10.1101/2025.04.07.647629

**Authors:** Avni Malhotra, Jessica A. M. Moore, Samantha Weintraub-Leff, Katerina Georgiou, Asmeret Asefaw Berhe, Sharon A. Billings, Marie-Anne de Graaff, Jennifer M Fraterrigo, A. Stuart Grandy, Emily Kyker-Snowman, Mingzhen Lu, Courtney Meier, Derek Pierson, Shersingh Joseph Tumber-Dávila, Kate Lajtha, William R Wieder, Robert B. Jackson

## Abstract

Increasing root carbon inputs into soils has been proposed as a solution to increasing soil organic carbon (SOC). However, while fine root carbon (FRC) inputs can increase SOC accrual in soils, FRC can also enhance SOC loss by stimulating microbial respiration and cause a net loss of SOC through priming. It remains unclear how SOC varies as a function of FRC at broad spatial scales and across ecosystems and depths. Here, we tested the relationship of SOC and FRC using data from 43 sites across the US National Ecological Observatory Network (NEON). We found that total stocks of SOC and FRC in the top 2 meters of soil were positively related with an across-ecosystem slope of 7 ± 3 kg SOC m^-2^ per kg FRC m^-2^. However, grassland sites primarily drove this relationship. Grasslands had 15 ± 2 kg SOC m^-2^ per kg FRC m^-2^, which is double the across-ecosystem slope. We used deviations from the standardized 1:1 relationship between FRC and SOC to infer whether ecosystems were net priming (indicated by observed SOC being lower than the 1:1 line) or SOC accruing (higher SOC than the 1:1 line). Grassland soils and especially their deep soil layers (>30 cm) showed primarily SOC accrual with increasing fine root abundance. Meanwhile, forest soils had high variability in whether increasing fine roots were associated with net SOC priming or accrual across both shallow and deeper soil layers. We found that in grasslands, FRC inputs are strongly related to SOC accrual, especially at depth and at sites with high moisture and clay content. In contrast, SOC-FRC relationships in forests remain difficult to characterize. Nevertheless, deep grassland soils may serve as optimal environments in which increasing FRC could lead to meaningful increases in SOC stocks.

## Introduction

Increasing and deepening root inputs into soils is proposed as a mechanism to increase soil organic carbon (SOC) but it remains unclear to what extent and under which environmental conditions this will be an effective strategy^1,2^. Fine roots (typically defined as roots with <2 mm diameter) are a key input to SOC and contribute disproportionately to SOC formation^3–5^. However, experimental evidence suggests that fine roots can either stabilize or destabilize SOC^6–9^. On one hand, labile compounds released by rhizodeposits or root litter may increase microbial biomass (and more critically, microbial necromass) thus increasing soil organic matter (SOM) if this necromass is stabilized by minerals^4,10,11^. On the other hand, the release of labile compounds from roots may cause a net loss of SOC (priming) by the breakdown of chemical associations between organic compounds and reactive soil minerals^12^, and by stimulating microbial respiration of detrital carbon (C)^13–15^.

Whether fine root C inputs drive SOC accrual or priming is expected to vary by ecosystem and vegetation type, soil moisture, SOC stock and its distribution between particulate and mineral-associated pools, the amount and reactivity of soil minerals, and macro and micro soil nutrients^6–8^. Since these factors vary throughout the organic and mineral layers of soils, we also expect soil depth and horizons to be important predictors of the relationship between fine root C and SOC. Ecosystem types with different dominant vegetation could also vary in their accrual and priming behaviors due to differences in belowground allocation, rooting depth and other root traits^16–18^. One emerging hypothesis is that SOC accrual is highest in soils with high reactivity minerals and in high moisture conditions, where plant productivity, SOM transport through the profile, and SOM stabilization to mineral surfaces are also high^13,19,20^. However, this hypothesis remains untested with regard to root-derived organic matter inputs. Quantifying the long-term stabilization of fine root C into SOC requires repeat measurements over multiple decades. Given the lack of such datasets^21–23^, natural gradients spanning variation in soil and ecosystem types, and containing soil and root measurements across depths, provide one means of testing the long-term, steady-state relationship between fine root C and SOC.

Here, we used a natural gradient with varying fine root biomass carbon stocks (hereafter, FRC) to explore the relationship between FRC and SOC stocks. We also tested how the FRC-SOC relationship varies by ecosystem type, soil depth and soil horizon (organic or mineral), and how the relationship is influenced by climate, mineralogy, and soil nutrients. We expected grassland ecosystem types to have stronger FRC-SOC relationships than forests because of the high below:aboveground biomass ratio in grasslands^17,18^. We also expected that mineral horizons would have a stronger relationship between FRC and SOC than organic horizons because the latter are likely more influenced by aboveground litter inputs^24^. Additionally, in mineral horizons where SOC stabilization can proceed via organo-mineral interactions, FRC is likely linked to net SOC to a greater extent than in organic horizons, where SOC may be less protected from microbial attack and microbial population abundances tend to be higher^5^.

Furthermore, by comparing our observed FRC-SOC relationships with a theoretical one-to-one relationship between FRC input and SOC stock, we inferred net SOC accrual versus priming across the gradient. Specifically, we assumed that sites with observed SOC above the 1:1 line of standardized SOC and FRC data indicate the potential for net SOC accrual (hereafter, SOC accrual), whereas sites with SOC below the 1:1 FRC-SOC line indicate the potential for net priming (hereafter, priming). We hypothesized that SOC accrual would be highest in ecosystems with high moisture and clay content where plant production and mineral stabilization of fine root litter would be optimized^13,19^. Conversely, priming would be more likely in ecosystems with lower moisture and clay content, where SOC would have a lower probability of interacting with soil minerals. We also hypothesized that SOC accrual would be greater at depth due to higher concentrations of reactive minerals and/or metals and lower microbial abundance/activity than in surface soils^25,26^.

We used the continental gradient provided by the National Ecological Observatory Network (NEON; Figure 1a) where coupled FRC and SOC measurements to 2-m depth have been conducted at 43 sites across the USA (see Table S1 for data sources). NEON sites represent a range of climates, with mean annual temperatures ranging from -12 to 25 °C and mean annual precipitation ranging from 100 to 2500 mm year^-1^. The sites also capture a variety of ecosystem types, though we focus our analyses primarily on grasslands and forests, which are the most abundant ecosystems across the network. Although it is difficult to leverage observational data collected at a continental spatial scale to probe processes such as microbial priming and mechanisms of SOC persistence that can occur at the micron scale, our work illuminates broad spatial patterns in SOC stocks that may be driven by these underlying processes.

**Figure 1.**
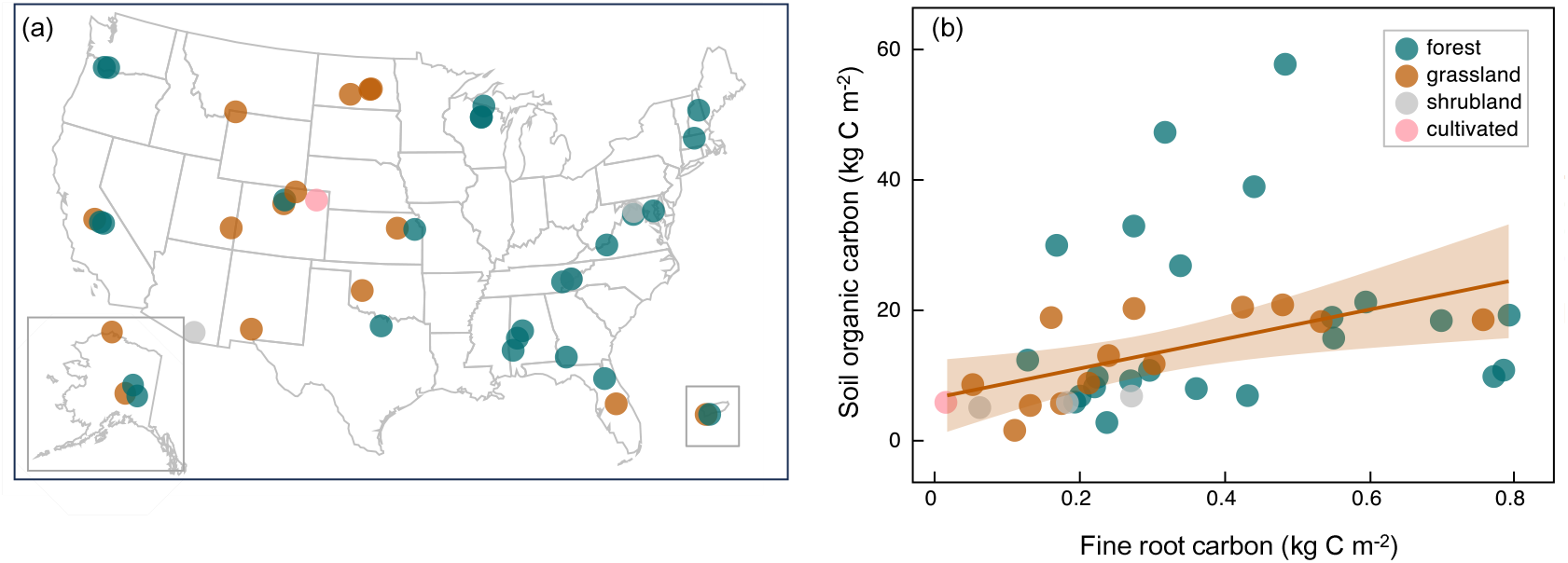
Site locations and relationships between fine root carbon and soil organic carbon stocks. (a) NEON site distribution by ecosystem type. Alaska and Puerto Rico are shown as insets. (b) The overall relationship between FRC and SOC (in Figure S1a) is driven by grasslands. Grasslands (n = 15) have a significant relationship between FRC and SOC (r^2^= 0.80, F_1,13_=53.14, p<0.0001; y = 8.5 + 15.5x). For ease of visualization, we have removed the three highest FRC sites here, but these tundra sites are included in Figure S1a and Table S2. Removing these outliers decreases the r^2^ to 0.46 but increases the slope to 22.6 kg SOC FRC^-1^ m^-2^. Forest FRC and SOC are unrelated (n = 25). Shrubland (n= 3) and cultivated (n= 1) ecosystem types do not have adequate sample sizes to analyze FRC-SOC relationships. Table S2 also provides information on depth of sampling, typically 2 m.

## Results

### Grasslands drive broad-scale fine root and soil carbon relationships

As hypothesized, FRC and SOC were related across sites, but these trends were primarily driven by grasslands (Figure 1b). Combining C stocks across the entire soil profile (to a maximum 2-m depth; Table S2), we found that FRC and SOC were positively related across our continental USA spatial gradient. The total SOC stock in a soil profile was positively related to whole-profile FRC (Figure S1a; adj r^2^= 0.39, p < 0.001, n = 43) and was best predicted by FRC, mean annual temperature (MAT), clay content, and land cover type (Table S3). For a one kg m^-2^ increase in FRC, there was a 7 ± 3 kg m^-2^ increase in SOC (Figure S1a). Within grasslands, for every one kg m^-2^ increase in FRC, there was a 15 ± 2 kg m^-2^ (p<0.0001 in a linear regression) increase in SOC (or 23 ± 7 kg m^-2^ increase in SOC if the two highest FRC outliers are excluded; p= 0.0114). This grassland-only estimate is more than double the value across all ecosystems (cross-ecosystem estimate: 7 ± 3 kg SOC m^-2^ per kg FRC m^-2^; Figure S1) due to the lack of a FRC-SOC relationship in forests. Our analyses suggest that aridity (MAP standardized by MAT; see methods), micronutrients, and aboveground litter input may be important in explaining forest SOC, but the relationships were not statistically significant (Figure S2).

When separated into organic and mineral horizons, FRC and SOC remained positively related across ecosystems, and ecosystem type was a significant predictor across most statistical models (Table S4a-b). However, the slope and best predictors of the relationship differed between soil horizons (Figure S1b). The only significant predictor of SOC in the organic horizon was FRC (adj r^2^= 0.41, p= 0.03, n= 17 out of which 2 were grasslands and rest were forests; Table S4a). Conversely, in the mineral horizon other factors such as MAT and percent clay were also important (adj r^2^= 0.30, p = 0.003, n= 43; Table S4b). Interestingly, three high latitude sites had more than twice as much root biomass than the others: NEON site codes WREF (cold and wet coniferous forest), BARR (tundra), and HEAL (tundra) (See Table S2 for details corresponding to site codes). Excluding these three sites, a model with root biomass, MAT, clay, and land cover still resulted in significant relationships, albeit weaker (adj r2 = 0.19, p < 0.001, n= 40). Thus, contrary to our hypotheses, our cross-ecosystem analysis suggests that in organic horizons, FRC is a primary predictor of SOC, while in mineral horizons, MAT and clay content are also important. MAT likely is a proxy for temperature limitations on plant productivity and decomposition of SOM while clay content represents the potential for mineral-associated organic matter formation.

### Strong positive FRC-SOC relationships in deep grassland soils

Similar to the relationships between total FRC and SOC summed across the soil profile, we found that depth distributions of FRC and SOC stocks (quantified using an exponential decay function fit; Figure S3) were related in grasslands but not in forests (Figure S3-S5, Tables S5a and S5b). Furthermore, we investigated how shallow (< 30 cm soil depth) and deep (> 30 cm soil depth) FRC influence shallow and deep SOC. Specifically, we tested whether the slope of the standardized FRC-SOC relationship in deep soil layers is higher than the slope of the FRC-SOC relationship in shallow layers. This observation would suggest a more effective SOC accrual per unit FRC at depth. We found that in deep soils, SOC increased more with increasing FRC than in shallow soils in grasslands (Figure S7 and Figure 2b) but not in forests (Figure S8 and Figure 2a). Our results thus support the hypothesis that deep FRC increases deep SOC more than shallow FRC increases shallow SOC, but only in grasslands. Furthermore, in grasslands, while a unit increase in standardized deep FRC increases deep SOC by 1.23 ± 0.42 (p =0.015), a unit increase in shallow FRC does not significantly increase shallow SOC (p= 0.19) (based on the slopes in Figure S7).

**Figure 2.**
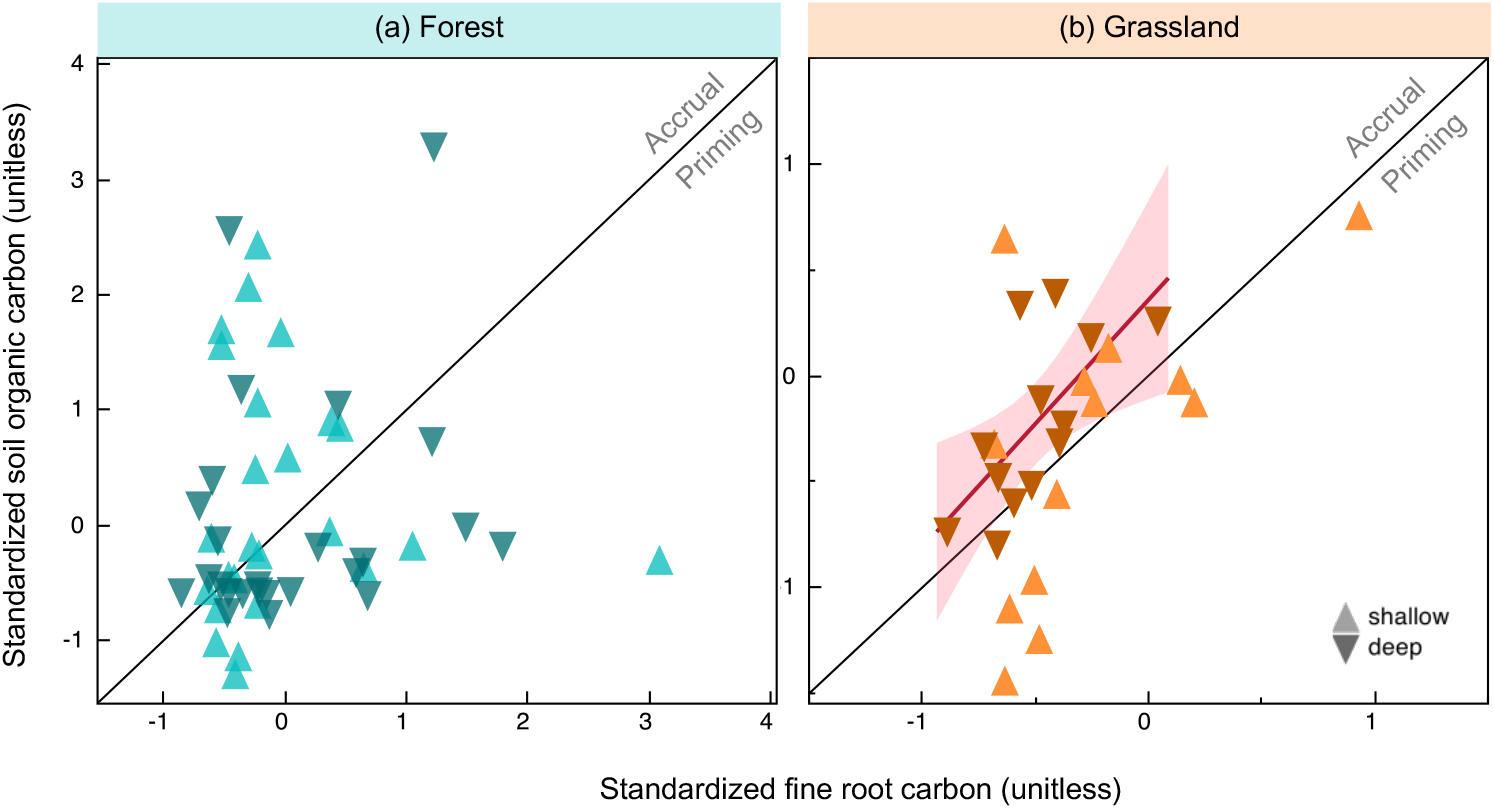
Standardized fine root carbon (FRC) and soil organic carbon (SOC). FRC and SOC are shown by soil depth along with a 1:1 line. Data points above the 1:1 line suggest inferred net accrual of SOC while data points below the 1:1 line suggest inferred priming. (a) Forests have no significant relationship between standardized FRC and SOC while (b) in grasslands the slope of the relationship between deep roots and deep SOC (>30 cm) is steeper (linear regression slope= 1.2, p=0.015) than the slope between shallow roots and shallow SOC (<30 cm depth; slope=0.9, p=0.19). Detailed statistics across depths and ecosystem type are provided in Figure S7 and S8.

### Lowest inferred priming in deep grassland soils

Using the standardized shallow and deep FRC-SOC relationships above, we quantified residuals from the 1:1 line, i.e., the difference between observed SOC value at a given site and the expected SOC value at a theoretical 1:1 line. We used these residuals as indicators of net SOC accrual or net priming relative to FRC inputs. Inferred SOC accrual corresponds to relatively more SOC being stored than incoming FRC (i.e., data points above the 1:1 line), and inferred priming corresponds to lower SOC being stored than incoming FRC (i.e., below the 1:1 line) (Figure 2). We found that forests exhibited great variation in inferred SOC accrual or priming relative to grasslands, which were primarily SOC accruing (Figure 3). In addition to ecosystem type, the degree of accrual or priming was predicted by factors related to moisture availability (aridity and MAP), soil texture and micronutrients (Table S6). In grasslands, priming increased with decreasing moisture availability, clay content and micronutrients; particularly in shallow soil layers (Figure S9). In forests, the variability in inferred priming was harder to explain, with the best-fit model explaining up to 44% of the variability in priming versus 73% in grasslands (Table S6). Deep forest soil dynamics remain particularly elusive as we could only explain up to 26% of the variability in inferred priming (Table S6). For example, we observed some indication of higher priming in warmer forests (Figure S10) compared to cooler forests, but this explained only 20% of the variability in priming.

**Figure 3.**
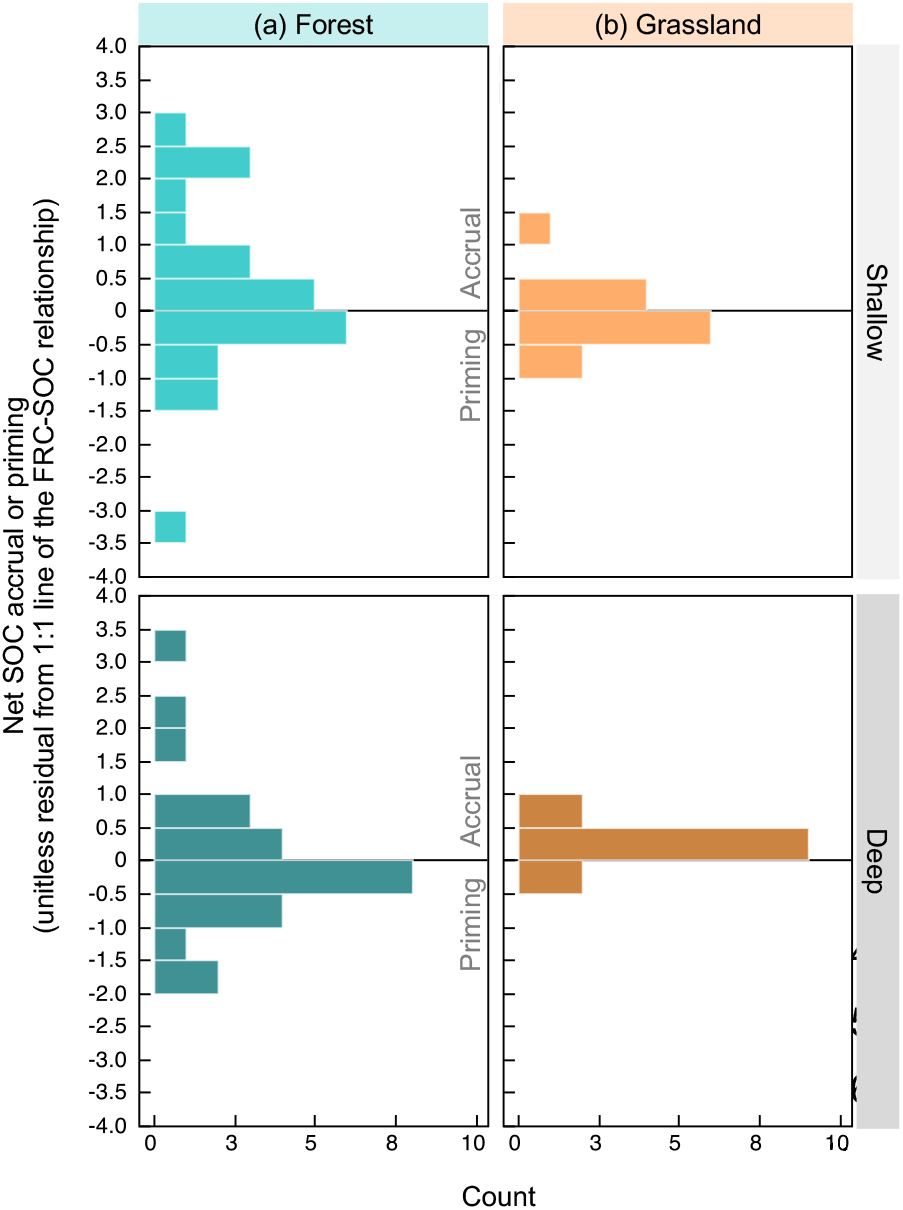
Distributions of inferred SOC accrual or priming. SOC accrual or priming is inferred from residuals of the 1:1 line between standardized FRC and SOC; see Figure 2. Data distributions are shown for shallow (<30 cm depth) and deep (30-200 cm) layers from both (a) forest and (b) grassland sites. Bins above the zero line are reflective of sites net SOC accrual and bins below the zero line reflect net priming. See Figure S12 for raw data points of the distributions.

## Discussion

### Unexplained variability in forest root-soil carbon relationships

There are several possible explanations for why forest FRC-SOC relationships and priming may be highly variable compared to grasslands. In this section, we use existing literature to discuss possible mechanisms behind our results.

First, grasslands typically have higher root:shoot ratios than forests and thus grassland root inputs may represent a dominant source of fresh carbon input into the soil^17,27,28^. Conversely, in forests, root:shoot can vary considerably across climate gradients and aboveground inputs may also be an important source of fresh carbon inputs into the soil, especially in deciduous forests^27^. This could mean that FRC inputs and FRC-induced priming are less influential on soil C processes compared to aboveground litter inputs in forests versus grasslands, although we did not see any evidence for aboveground litterfall rates predicting forest SOC either (Figure S2). It is possible that increased aboveground litterfall also led to priming in forests, as previously observed in temperate and tropical forests^29–32^.

Second, in forests and in woody plants, maximum rooting depths vary widely and differ from the more conserved rooting depths of grasslands^16,33^. This rooting depth variation would also influence the relationship between FRC and SOC, given the hypothesis that deeper roots have a greater propensity for mineral interactions and thus contribute to persistent SOC pools^26^. In our data we also observed generally deeper and more variable depth distributions of FRC and SOC in forests compared to grasslands (Figure S4). The variability in forest FRC-SOC relationships suggests that factors other than FRC contribute to forest SOC accrual to a greater extent than in grassland soils.

Third, the chemical composition of litter and root inputs may vary more across forest types compared to grasslands, making the relationship between FRC inputs and SOC accumulation across forests harder to predict. Graminoid species can be chemically relatively simple compared to other plants^34^. In contrast, tree species are known to differentially affect decomposition and soil C and N cycling^35,36^, including the magnitude of the rhizosphere priming effect^37^. Leaf and root litter from some tree species can be a source of tannins and other phenolics that affect soil processes^38–40^, and the presence or absence of tannin-rich species on our forest sites may represent an important source of variability in forest SOC dynamics. For example, tannin-rich litter could promote SOC stabilization while cellulose-rich litter could stimulate priming^41^. Thus, plant litter quality influences priming or stabilization of SOC, and greater variability in litter quality in forests compared to grasslands could potentially contribute to our observed high variation in forest SOC dynamics^42^.

Fourth, grassland plant roots typically have higher absorptive:transport root ratios and a greater proportion of absorptive roots with short lifespans, which could imply greater exudation and greater root litter contribution to SOC, respectively^43,44^. Given the generally greater physiological activities of fine compared to coarse roots^43^, a greater relative abundance of high-turnover, absorptive roots in grasslands compared to forests^45^ might also result in greater activity rates of soil microbes. This greater microbial activity could also be higher in grasslands than in forests because grassland roots decompose faster than forest roots^46^. Thus, we might expect greater rates of microbial necromass production where fine root abundance is relatively greater, given fine root exudate activities and associated rhizosphere microbial growth and death^47^. Evidence is accumulating that microbial necromass is a meaningful component of persistent SOC stocks^47–49^. We might expect, then, that where absorptive root litter production is greater, necromass and thus SOC accrual might also be greater.

Fifth, higher and more variable precipitation in forests compared to grasslands could drive variability in the activity of decomposition enzymes and reactive metals^19^. Thus, mineralogical limits to C storage, driven by variations in the amount and reactivity of clay minerals and reactive metals, may be more important controls on forest SOC than litter inputs^19,50^. Since our predictive models do not include detailed proxies defining mineralogical limits to C storage, we may be missing some explanatory power and thus seeing high variability in forests. Furthermore, our grassland sites, on average, have half the mean annual precipitation of our forest sites. Thus, plant growth, litter inputs and decomposition could all be more moisture limited in grasslands than in forests, such that C inputs and not mineralogical limits are a more important factor for stabilizing SOC in grasslands.

Lastly, forests have the added complexity of a variety of mycorrhizal symbionts that could be influencing plant-soil processes and belowground carbon allocation in ways different than in grasslands, which are often limited to arbuscular mycorrhizal types that have a lower carbon demand (Figure S11)^51–53^. Dominant mycorrhizal association type was not a significant predictor in any of our models, but it is likely that fungal biomass would have been a better predictor (data unavailable). FRC and fungal biomass together would better capture the forest variability in total belowground carbon allocation. Furthermore, high variation in mycorrhizal types and other microbial community types in forests would also lead to diverse decomposition dynamics^e.g., 54^. Overall, it makes sense that while we were able to explain inferred priming in grasslands using simple climate, soil texture, and nutrient information, inferred priming in forests would require additional predictors including information about root systems, litter quality, mineralogical limits to C storage, and microbial community structure and function.

### Limitations

Our inferred priming proxy allows us to explore potential relationships between FRC and SOC and offers hypotheses to probe the drivers of SOC formation and persistence. However, our approach has limitations. First, our proxy does not account for all the belowground C inputs into SOC. We only consider standing fine root biomass in our calculations. Ideally, inputs to SOC should include rhizodeposits, and should incorporate information about root and fungal turnover rates^55^, but these data were unavailable. Second, we did not have a way to account for variation in the decomposition of incoming root litter. In other words, at sites where we saw high inferred priming, there may have simply been efficient decomposition of fresh root litter. Root litter decomposition rates in the first year can vary widely in forests, with estimates suggesting 20-40% mass loss^56^. Thus, we expect that this variability further contributed to the lack of a clear FRC-SOC relationship in forests. Third, as is the case with many observational studies, it is difficult to ascribe causation to correlative relationships. A variety of climate and edaphic factors could be driving both FRC and SOC, thus resulting in the observed relationships.

## Conclusions

In the last few decades, paradigm shifts in SOC research suggest that root carbon inputs are central to organic matter formation and stabilization^5^. We found that at broad spatial scales, FRC and SOC are related but these trends were driven by grasslands and not forests across the NEON gradient. We also investigated whether deeper roots are associated with higher deep SOC, which is presumed to be more stable, and found support for this in grasslands but again not in forests. Our hypothesis that stabilization of FRC into SOC would be highest under conditions of high moisture and mineral surface availability (using clay content as a proxy) was supported in grasslands. Future data collection efforts at the continental scale and beyond should quantify other belowground carbon inputs (e.g., root turnover and exudation, and microbial biomass turnover) into soils to enable a more mechanistic understanding of FRC and SOC linkages across biomes. Nevertheless, in the context of management strategies of increasing FRC to increase SOC, root biomass will likely be the main trait that can be measured and managed, and not root turnover^2^. Thus, our analysis provides a useful benchmark of how FRC and SOC are related across broad scales and ecosystems.

Unlike grasslands, forests displayed high variability in FRC-SOC relationships, both across space and depth, likely due to the potentially higher complexity in root-soil interactions in forests. Thus, forests may not be ideal settings to increase SOC through fresh root carbon inputs as this may result in priming-induced SOC losses and increased CO_2_ emissions via increased root litter and SOM decomposition.

Predictors of this forest priming effect, especially in deeper layers, remain elusive and can serve as an important future research trajectory. Grassland soils with relatively high moisture and clay content, may serve as settings in which increased root-derived SOC sequestration may be promoted at depth, although replication across climate gradients and detailed measurements of root dynamics are needed to confidently project SOC accrual.

## Materials and methods

### Sites and data

The root and soil carbon data used for this study were collected by NEON, a continental-scale ecological monitoring program spanning 47 terrestrial sites and all major US ecoclimatic regions. We used data from the NEON ‘megapits’, which include measurements of soil chemical properties, physical properties, and root biomass (Table S1). We downloaded these data from the Soils Data Harmonization (SoDaH) database ^22^ (data accessed Jul 2020) which includes NEON data^57^ among other network data sources. Data from other networks were not considered in this study because sites infrequently measured root biomass and soil chemistry profiles to depth in the same location. Four NEON sites (STER, KONA, PUUM, and TOOL) were excluded from our study because root data were not collected in the NEON ‘megapits’.

Thus, 43 of the 47 NEON sites were included in our analyses. All 43 sites had mineral soil horizons, and 17 of the 43 sites had organic soil horizons (15 forests and 2 high latitude grasslands/tundra). Site metadata such as mean annual temperature, mean annual precipitation, and site-wide dominant plants were taken from SanClements et al ^58^. We also calculated an aridity index as MAP (mm) standardized by MAT (degree Celsius) using the formula MAP/(MAT+13). The 13 was added to adjust for negative MAT values ^59^ and lower values represent more arid conditions. Land cover was ascertained by NEON scientists and was based on the NLCD land cover classifications ^60^ from NEON field site information tables ([link](https://www.neonscience.org/field-sites/explore-field-sites)).

### Sample collection and processing

The NEON megapit sampling effort was a one-time measurement conducted by NEON staff and the USDA Natural Resource Conservation Service (NRCS) over the course of 2014-2018. At each site, NEON scientists and contractors excavated a 2 m deep (or to bedrock) soil pit in the vicinity of the NEON eddy covariance tower. The timing of sampling varied across the growing season and was not always at peak biomass. NRCS soil scientists then assigned soil taxonomy *in situ*, and by taxonomic horizon to the bottom of the pit. These samples were then sent to the Kellogg Soil Survey Laboratory in Lincoln, Nebraska where, after passing through a 2 mm sieve, they were analyzed for a host of physical and chemical properties including bulk density, particle size, total C, nitrogen (N), phosphorous (P), metals, and other edaphic properties using standard NRCS methods ^61^.

At each NEON megapit, root samples were collected across depth profiles. Samples were collected in 10-cm depth increments to 1 m depth, then in 20-cm depth increments to 2 m depth by cutting 10-cm deep x 10 cm wide soil monoliths, in three vertical profiles on the left, middle, and right side of the pit. Roots were hand-sorted from these monoliths, visually classified as live or dead, and diameter was measured. Most NEON sites (30 sites out of 43) classified “fine roots” as less than 2 mm diameter. However, 13 sites had a different methodological protocol and used a 4 mm diameter cutoff. Of these, nine were forests, three were rangelands/grasslands (hereafter, grassland), and one was a shrubland. Note that we statistically tested the hypothesis that having different fine root diameter cut offs may be related to our observed variability in forest FRC-SOC relationships. This hypothesis was not supported in a Wilcoxon rank test of the residuals from the FRC-SOC relationship when comparing the two diameter classes (Z= 0.59, p= 0.55). Other studies^62^ have also found that root biomass from these two diameter classes are highly correlated across sites in fine root databases, and that in the NEON sites, the diameter sampling differences do not influence properties such as rooting depth distribution^62^.

Root biomass was measured after oven-drying the samples at 65°C for at least 48 hours. Dried root samples were sent to the University of Wyoming for analysis of C concentrations using elemental analysis. The three vertical pit profiles per megapit were averaged prior to ingestion into SoDaH and used in our statistical analysis. Despite the 4 mm diameter exceptions, we consider the root stock to represent ‘fine-root biomass’ throughout the manuscript.

Lastly, we used annual litterfall fluxes (forest sites only) as a covariate in an exploratory analysis (see Table S1 for data sources). Briefly, annual litterfall was measured by collecting all material dropped from the forest canopy with a diameter <2 cm and a length <50 cm using elevated 0.5 m^2^ PVC traps. Traps were deployed (20 plots per site) near the megapits. We used the total mass (leaves, needles, twigs, etc. all added) collected by the traps over the course of a growing season to estimate annual productivity. Where multiple years of data were available, the average flux was used.

### Data alignment

Alignment of FRC and SOC data was necessary due to different sampling strategies for roots and soils. Roots were sampled at fixed (10 or 20 cm) increments through the profile, while soils were sampled once in each taxonomic horizon regardless of horizon depth. Therefore, we aligned the root data with the corresponding soil horizon. Fine root biomass C stocks (FRC) were calculated as the product of root biomass (g m^-2^) in a given depth interval and FRC concentration (%). SOC stocks were calculated as the product of soil organic C concentration (%), bulk density (kg/cm^3^), and sampling depth (cm), then converted to kg m^-2^ by multiplying by 1 × 10^3^.

### Calculation of beta coefficients

In order to investigate the relationship between depth profiles of FRC and SOC, we calculated beta values using an exponential decay curve (Eq. 1), which describes how stocks change with depth ^18,63^. Of the 43 NEON sites, 36 were used to calculate beta coefficients. Seven sites (BARR, CLBJ, GRSM, GUAN, JORN, LAJA, TEAK) were excluded because beta coefficients could not be calculated due to too few SOC measurements in the profile. Soil organic carbon and FRC may accumulate primarily in surface soils and to varying degrees deeper in the profile, or there may be a gradual and consistent increase at each depth interval. These different accumulation patterns can be captured in an exponential function (Eq. 1), where a higher beta coefficient indicates a deeper distribution of root or SOC, relative to a lower beta. We converted each depth profile (FRC or SOC) at each site into one beta coefficient to facilitate these analyses.

Beta coefficients (*β*) were calculated using the following function ^18,63^:

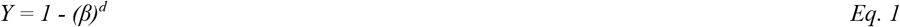

In Eq. 1, *Y* is the cumulative fraction of either SOC or root biomass in a given layer with respect to the whole profile, and *d* is the depth (cm) measured at the bottom of that layer. For every depth layer at every site, we solved for *β* in Eq. 1 for both SOC (*β*_*SOC*_) and root biomass (*β*_*roots*_). These *β* values were used as starting parameters at discrete points through the depth profile, and we used the iterative Bound Approximation by Quadratic Approximation (BOBYQA) method (package minqa in R) ^64^ to interpolate between points and resolve the function across the continuous depth profile at each site ^65^. The BOBYQA-resolved *β*_*SOC*_ and *β*_*FRC*_ values were used as response and fixed effect variables, respectively, in mixed effects models.

### Statistical analyses

We analyzed relationships between SOC, FRC, and other climatic and edaphic covariates using linear mixed-effects models. We analyzed FRC-SOC relationships in three ways wherein FRC and SOC stocks were: 1) summed across the whole profile, 2) separated by organic and mineral soil horizons, and 3) described as beta coefficients as a function of depth. For each analysis, we constructed a null (random effects only) model, a full model, and then reduced models that lacked covariates. We selected best-fit models based on the lowest Akaike Information Criterion (AIC) score aside from the full model, to avoid overparameterization, or the sample-size corrected AIC (AICc) score when the data set contained fewer than 40 observations ^66^. In whole-profile and by-horizon analyses, the full model included SOC as the response and FRC, mean annual temperature (MAT), mean annual precipitation (MAP), clay percent, and land cover (ecosystem type) as fixed effects and maximum profile depth as a random effect. We also verified that sampling depth does not influence our analyses by conducting a multiple regression analysis including all covariates and maximum profile depth and found that profile depth was not a significant parameter (p = 0.57, partial r^2^ = 0.01). Lastly, we explored the role of mycorrhizal associations by adding dominant mycorrhizal type (arbuscular, ectomycorrhizal or mixed) ^51^ as a fixed effect but saw no significant relationships or model improvements. Across the mixed-effects models, we report the significance level (p-value) calculated using Satterthwaite’s method (lmerTest R package) ^67^, a test statistic (χ^2^), and marginal pseudo-R^2^ (sjstats R package)^68^. The fixed effects of the best fit model were tested using analysis of variance (Anova function in the R package car)^69^. Forest and grassland land cover types were tested and shrublands and cultivated lands were excluded from this analysis due to limited sample size. Assumptions of homoscedasticity, low variance inflation factors and normal data distributions were verified for each statistical model.

### Inferred SOC accrual and priming

We calculated inferred net accrual or priming using residuals, which measure the difference between observed and predicted values, from the shallow (<30 cm) and deep (>30 cm depth) FRC-SOC relationships. Specifically, we determined the residual difference between the observed SOC and expected SOC along a standardized 1:1 line, which represents a theoretical scenario where each unit of FRC input results in an equivalent unit increase in SOC (Figure S7 and S8). This 1:1 line serves as the baseline for assessing if SOC levels are higher or lower than expected based on FRC inputs. The calculated residuals, therefore, act as a proxy for SOC accrual (when observed values exceed the expected) or priming (when observed values are less than expected) in relation to FRC inputs. Furthermore, we employed multiple regression models to explore the potential factors influencing this inferred priming or accrual, as detailed in Table S6. These models allow us to identify and evaluate variables that may affect the relationship between FRC and SOC across different sites and conditions.

## Acknowledgements

AM was supported by the Gordon and Betty Moore Foundation (Grant GBMF5439) and the Laboratory Directed Research and Development (LDRD) Program at Pacific Northwest National Laboratory under U.S. Department of Energy (DOE) contract DE-AC05-76RL01830. This work was also supported by the Long Term Ecological Research Network Office (LNO; NSF awards 1545288 and 1929393) and the National Center for Ecological Analysis and Synthesis at the University of California Santa Barbara (Awarded to KL and WRW). The National Ecological Observatory Network (NEON) is a program sponsored by the U.S. National Science Foundation and operated under cooperative agreement by Battelle. This material is based in part upon work supported by the U.S. National Science Foundation through the NEON Program. KG was supported by the LLNL-LDRD Program under the auspices of DOE Contract DE-AC52-07NA27344. SJTD acknowledges support from the US National Science Foundation (NSF-DEB-LTER 1832210). WRW acknowledges support from NSF award numbers (2224439 and 1926413).

## Author contributions

AM, JAM, SW-L, KG, EK-S, ASG, DP, KL, WRW, RBJ, SJT-D and ML conceptualized the study. AM, JAM, SW-L and KG performed the analyses. AAB, SAB, MG, JMF, ASG, CM, DP, KL, WRW, and SW-L provided data. AM wrote the manuscript with input from all coauthors. RBJ, KL and WRW acquired funding.

## Data availability

All data used in this manuscript are publicly available from the National Ecological Observatory Network (NEON) and the corresponding data product names are reported in Supplementary Table S1.

## Code availability

Code used in the study is available as a supplementary materials file.

## References

1. Bossio, D. A. et al. The role of soil carbon in natural climate solutions. Nature Sustainability 3, 391– 398 (2020).

2. Kell, D. B. Breeding crop plants with deep roots: their role in sustainable carbon, nutrient and water sequestration. Ann. Bot. 108, 407–418 (2011).

3. Jackson, R. B. et al. The Ecology of Soil Carbon: Pools, Vulnerabilities, and Biotic and Abiotic Controls. Annu. Rev. Ecol. Evol. Syst. 48, 419–445 (2017).

4. Rasse, D. P., Rumpel, C. & Dignac, M.-F. Is soil carbon mostly root carbon? Mechanisms for a specific stabilisation. Plant Soil 269, 341–356 (2005).

5. Rocci, K. S. et al. Bridging 20 years of soil organic matter frameworks: Empirical support, model representation, and next steps. J. Geophys. Res. Biogeosci. 129, e2023JG007964 (2024).

6. Lajtha, K., Bowden, R. D. & Nadelhoffer, K. Litter and root manipulations provide insights into soil organic matter dynamics and stability. Soil Sci. Soc. Am. J. 78, S261–S269 (2014).

7. Bowden, R. D. et al. Litter Input Controls on Soil Carbon in a Temperate Deciduous Forest. Soil Science Society of America Journal 78, S66–S75 (2014).

8. Pierson, D. et al. Mineral stabilization of soil carbon is suppressed by live roots, outweighing influences from litter quality or quantity. Biogeochemistry 154, 433–449 (2021).

9. Dijkstra, F. A., Zhu, B. & Cheng, W. Root effects on soil organic carbon: a double-edged sword. New Phytol. 230, 60–65 (2021).

10. Sokol, N. W. et al. The path from root input to mineral-associated soil carbon is dictated by habitat-specific microbial traits and soil moisture. Soil Biol. Biochem. 193, 109367 (2024).

11. Miltner, A., Bombach, P., Schmidt-Brücken, B. & Kästner, M. SOM genesis: microbial biomass as a significant source. Biogeochemistry 111, 41–55 (2012).

12. Keiluweit, M. et al. Mineral protection of soil carbon counteracted by root exudates. Nat. Clim. Chang. 5, 588–595 (2015).

13. Bastida, F. et al. Global ecological predictors of the soil priming effect. Nat. Commun. 10, 3481 (2019).

14. Bailey, V. L., Pries, C. H. & Lajtha, K. What do we know about soil carbon destabilization? Environ. Res. Lett. 14, 083004 (2019).

15. Kuzyakov, Y., Friedel, J. K. & Stahr, K. Review of mechanisms and quantification of priming effects. Soil Biol. Biochem. 32, 1485–1498 (2000).

16. Tumber-Dávila, S. J., Schenk, H. J., Du, E. & Jackson, R. B. Plant sizes and shapes above and belowground and their interactions with climate. New Phytol. 235, 1032–1056 (2022).

17. Qi, Y., Wei, W., Chen, C. & Chen, L. Plant root-shoot biomass allocation over diverse biomes: A global synthesis. Global Ecology and Conservation 18, e00606 (2019).

18. Jackson, R. B. et al. A global analysis of root distributions for terrestrial biomes. Oecologia 108, 389–411 (1996).

19. Heckman, K. A. et al. Moisture-driven divergence in mineral-associated soil carbon persistence. Proc. Natl. Acad. Sci. U. S. A. 120, e2210044120 (2023).

20. Cusack, D. F. & Turner, B. L. Fine Root and Soil Organic Carbon Depth Distributions are Inversely Related Across Fertility and Rainfall Gradients in Lowland Tropical Forests. Ecosystems 24, 1075– 1092 (2021).

21. Malhotra, A. et al. The landscape of soil carbon data: Emerging questions, synergies and databases. Progress in Physical Geography: Earth and Environment 43, 707–719 (2019).

22. Wieder, W. R. et al. SoDaH: the SOils DAta Harmonization database, an open-source synthesis of soil data from research networks, version 1.0. Earth System Science Data 13, 1843–1854 (2021).

23. Weintraub, S. R. et al. Leveraging Environmental Research and Observation Networks to Advance Soil Carbon Science. Journal of Geophysical Research: Biogeosciences 124, 1047–1055 (2019).

24. Garten, C. T. A disconnect between O horizon and mineral soil carbon – Implications for soil C sequestration. Acta Oecologica 35, 218–226 (2009).

25. Fraterrigo, J. M., Ream, K. & Knoepp, J. D. Tree Mortality From Insect Infestation Enhances Carbon Stabilization in Southern Appalachian Forest Soils. Journal of Geophysical Research: Biogeosciences 123, 2121–2134 (2018).

26. Rumpel, C. & Kögel-Knabner, I. Deep soil organic matter—a key but poorly understood component of terrestrial C cycle. Plant Soil 338, 143–158 (2011).

27. Ma, H. et al. The global distribution and environmental drivers of aboveground versus belowground plant biomass. Nat Ecol Evol 5, 1110–1122 (2021).

28. Steinaker, D. F. & Wilson, S. D. Belowground litter contributions to nitrogen cycling at a northern grassland–forest boundary. Ecology 86, 2825–2833 (2005).

29. Sayer, E. J., Heard, M. S., Grant, H. K., Marthews, T. R. & Tanner, E. V. J. Soil carbon release enhanced by increased tropical forest litterfall. Nat. Clim. Chang. 1, 304–307 (2011).

30. Crow, S. E. et al. Increased coniferous needle inputs accelerate decomposition of soil carbon in an old-growth forest. For. Ecol. Manage. 258, 2224–2232 (2009).

31. Fontaine, S. et al. Stability of organic carbon in deep soil layers controlled by fresh carbon supply. Nature 450, 277–280 (2007).

32. Sayer, E. J. et al. Tropical forest soil carbon stocks do not increase despite 15 years of doubled litter inputs. Sci. Rep. 9, 18030 (2019).

33. Canadell, J. et al. Maximum rooting depth of vegetation types at the global scale. Oecologia 108, 583–595 (1996).

34. Meier, C. L. & Bowman, W. D. Links between plant litter chemistry, species diversity, and below-ground ecosystem function. Proc. Natl. Acad. Sci. U. S. A. 105, 19780–19785 (2008).

35. Raich, J. W. & Tufekciogul, A. Vegetation and soil respiration: Correlations and controls. Biogeochemistry 48, 71–90 (2000).

36. Hobbie, S. E. et al. Tree species effects on decomposition and forest floor dynamics in a common garden. Ecology 87, 2288–2297 (2006).

37. Yin, L., Dijkstra, F. A., Wang, P., Zhu, B. & Cheng, W. Rhizosphere priming effects on soil carbon and nitrogen dynamics among tree species with and without intraspecific competition. New Phytol. 218, 1036–1048 (2018).

38. Fierer, N., Schimel, J. P., Cates, R. G. & Zou, J. Influence of balsam poplar tannin fractions on carbon and nitrogen dynamics in Alaskan taiga floodplain soils. Soil Biol. Biochem. 33, 1827–1839 (2001).

39. Talbot, J. M. & Finzi, A. C. Differential effects of sugar maple, red oak, and hemlock tannins on carbon and nitrogen cycling in temperate forest soils. Oecologia 155, 583–592 (2008).

40. Adamczyk, B. et al. Plant roots increase both decomposition and stable organic matter formation in boreal forest soil. Nat. Commun. 10, 3982 (2019).

41. Huys, R. et al. Plant litter chemistry controls coarse-textured soil carbon dynamics. J. Ecol. 110, 2911–2928 (2022).

42. Craig, M. E. et al. Fast-decaying plant litter enhances soil carbon in temperate forests but not through microbial physiological traits. Nat. Commun. 13, 1229 (2022).

43. McCormack, M. L. et al. Redefining fine roots improves understanding of below-ground contributions to terrestrial biosphere processes. New Phytol. 207, 505–518 (2015).

44. Chari, N. R. et al. Estimating the global root exudate carbon flux. Biogeochemistry 167, 895–908 (2024).

45. Jackson, R. B., Mooney, H. A. & Schulze, E. D. A global budget for fine root biomass, surface area, and nutrient contents. Proc Natl Acad Sci U S A 94, 7362–7366 (1997).

46. Solly, E. F. et al. Factors controlling decomposition rates of fine root litter in temperate forests and grasslands. Plant Soil 382, 203–218 (2014).

47. Sokol, N. W. et al. Life and death in the soil microbiome: how ecological processes influence biogeochemistry. Nat. Rev. Microbiol. 20, 415–430 (2022).

48. Simpson, A. J., Simpson, M. J., Smith, E. & Kelleher, B. P. Microbially derived inputs to soil organic matter: are current estimates too low? Environ Sci Technol 41, 8070–8076 (2007).

49. Liang, C., Amelung, W., Lehmann, J. & Kästner, M. Quantitative assessment of microbial necromass contribution to soil organic matter. Glob Chang Biol 25, 3578–3590 (2019).

50. Rasmussen, C. et al. Beyond clay: towards an improved set of variables for predicting soil organic matter content. Biogeochemistry 137, 297–306 (2018).

51. Chaudhary, V. B. et al. MycoDB, a global database of plant response to mycorrhizal fungi. Sci Data 3, 160028 (2016).

52. Brzostek, E. R., Fisher, J. B. & Phillips, R. P. Modeling the carbon cost of plant nitrogen acquisition: Mycorrhizal trade-offs and multipath resistance uptake improve predictions of retranslocation. J. Geophys. Res. Biogeosci. 119, 1684–1697 (2014).

53. Yang, K. et al. Mycorrhizal type regulates trade-offs between plant and soil carbon in forests. Nat. Clim. Chang. 14, 91–97 (2023).

54. Xu, Z. et al. The variations in soil microbial communities, enzyme activities and their relationships with soil organic matter decomposition along the northern slope of Changbai Mountain. Appl. Soil Ecol. 86, 19–29 (2015).

55. Poirier, V., Roumet, C. & Munson, A. D. The root of the matter: Linking root traits and soil organic matter stabilization processes. Soil Biol. Biochem. 120, 246–259 (2018).

56. Berg, B., Johansson, M., Meentemeyer, V. & Kratz, W. Decomposition of tree root litter in a climatic transect of coniferous forests in northern Europe: A synthesis. Scand. J. For. Res. 13, 402– 412 (1998).

57. NEON (National Ecological Observatory Network). Soil physical and chemical properties, Megapit (DP1.00096.001, accessed January 1, 2020); Root biomass and chemistry, Megapit (DP1.10096.001, accessed January 1, 2020); and Litterfall and fine woody debris production and chemistry (DP1.10033.001, accessed January 1 2021); PROVISIONAL. Data accessed from https://data.neonscience.org/.

58. SanClements, M. et al. Collaborating with NEON. BioScience 70, 107–107 (2020).

59. Croitoru, A.-E., Piticar, A., Imbroane, A. M. & Burada, D. C. Spatiotemporal distribution of aridity indices based on temperature and precipitation in the extra-Carpathian regions of Romania. Theor. Appl. Climatol. 112, 597–607 (2013).

60. Homer, C. H., Fry, J. A. & Barnes, C. A. The national land cover database. US Geological Survey Fact Sheet 3020, 1–4 (2012).

61. Burt, R. Soil Survey Staff. Kellogg Soil Survey Laboratory methods manual. US Department of Agriculture, Natural Resources Conservation Service, National Soil Survey Center, Kellogg Soil Survey Laboratory, Lincoln, NE (2014).

62. Lu, M. et al. A continental scale analysis reveals widespread root bimodality. bioRxiv (2022) doi:10.1101/2022.09.14.507823.

63. Gale, M. R. & Grigal, D. F. Vertical root distributions of northern tree species in relation to successional status. Canadian Journal of Forest Research 17, 829–834 (1987).

64. Mullen, K. M. Minqa: Derivative-Free Optimization Algorithms by Quadratic Approximation. (2024).

65. Powell, M. The BOBYQA algorithm for bound constrained optimization without derivatives. (2009).

66. Wagenmakers, E.-J. & Farrell, S. AIC model selection using Akaike weights. Psychon. Bull. Rev. 11, 192–196 (2004).

67. Kuznetsova, A., Brockhoff, P. B. & Christensen, R. H. B. lmerTest Package: Tests in Linear Mixed Effects Models. J. Stat. Softw. 82, 1–26 (2017).

68. Lüdecke, D. Sjstats: Statistical Functions for Regression Models. doi:10.5281/zenodo.1489175.

69. Fox, J. An R Companion to Applied Regression. (2019).

