## Supplementary Information for "Continental-scale relationships of fine root and soil carbon stocks hold in grasslands but not forests"

[1]Department of Earth System Science, Stanford University, Stanford, CA, USA;; 0000-0002-7850-6402
*Current address: Biological Sciences Division, Pacific Northwest National Laboratory, Richland, WA, USA
[3]National Ecological Observatory Network, Battelle, Boulder CO USA;, 0000-0003-4789-5086
, 0000-0003-3576-883X
[4]Department of Biological and Ecological Engineering, Oregon State University, Corvallis, OR, USA;, 0000-0002-2819-3292
[5]Department of Life and Environmental Sciences; University of California, Merced, Merced, CA, USA;; 0000-0002-6986-7943
[6]Department of Ecology and Evolutionary Biology, Kansas Biological Survey & Center for Ecological Research, University of Kansas, Lawrence, KS, USA;; 0000-0003-1611-526X
[7]Department of Biological Sciences, Boise State University, Boise, ID, USA;; 0000-0001-5668-7647
[8]Department of Natural Resources and Environmental Sciences, University of Illinois, Urbana, IL, 61801, USA;; 0000-0002-0357-1007
[9]Department of Natural Resources and the Environment, University of New Hampshire, Durham, NH, USA
[10]Carbon Direct, New York, NY, USA; 0000-0003-1782-1916
[11] New York University, New York, NY, USA;; 0000-0002-8707-8745
[12]Rocky Mountain Research Station, United States Forest Service, Boise, ID, USA; ; 0000-0003-3413-1693
[13]Department of Environmental Studies, Dartmouth College, Hanover, NH, USA;; 0000-0001-7336-3943
[14]Department of Crop and Soil Science, Oregon State University, Corvallis, OR, USA; 0000-0002-6430-4818
[15]Climate and Global Dynamics Laboratory, National Center for Atmospheric Research, Boulder, CO 80307, USA & Institute of Arctic and Alpine Research, University of Colorado, Boulder, CO 80309, USA;; 0000-0001-7116-1985
[16]Woods Institute for the Environment and Precourt Institute for Energy, Stanford University, Stanford, CA 94305

Table S1. NEON data products used in the analysis.

| Data Product ID | Product Name | Variables | Date Range |
| --- | --- | --- | --- |
| DP1.00096.001 | Soil physical and chemical properties, Megapit | Particle size distribution, bulk density, soil C, nutrient and metal concentrations | 2014-2018 |
| DP1.10066.001 | Root biomass and chemistry, Megapit | Root biomass and C concentration | 2014-2018 |
| DP1.10033.001 | Litterfall and fine woody debris production and chemistry (forest sites only) | Litterfall fluxes | 2016-2019 |

Table S2. Fine-root biomass, SOC and betas of fine roots and SOC for the full profile (i.e., organic and mineral horizons) at each NEON site where it was possible to calculate beta coefficients (n = 37). Note that a handful of profiles did not have 200 cm as their bottom depth (due to sampling or bedrock differences). Also, STER was removed from the mixed models that use land cover because it is the only one representing ‘cultivated’.

| NEON site code | Land cover | Ecoregion | Maximum measurement depth (cm) | Fine-root biomass (kg C $m^{-2}$) | SOC (kg C $m^{-2}$) |
|---|---|---|---|---|---|
| BART | forest | temperate broadleaf or mixed forest | 200 | 0.54 | 16.11 |
| HARV | forest | temperate broadleaf or mixed forest | 160 | 0.78 | 19.51 |
| RMNP | forest | temperate coniferous forest | 160 | 0.29 | 30.83 |
| CPER | grassland | temperate grassland, savanna and shrubland | 200 | 0.21 | 9.02 |
| STER | cultivated | temperate grassland, savanna and shrubland | 200 | 0.01 | 5.71 |
| OAES | grassland | temperate grassland, savanna and shrubland | 46 | 0.05 | 8.85 |
| YELL | grassland | temperate grassland, savanna and shrubland | 200 | 0.43 | 22.24 |
| MOAB | grassland | desert or xeric shrubland | 200 | 0.17 | 5.86 |
| NIWO | grassland | tundra | 200 | 0.76 | 18.84 |
| ONAQ | shrubland | desert or xeric shrubland | 200 | 0.27 | 7.06 |
| BLAN | shrubland | temperate grassland, savanna and shrubland | 200 | 0.18 | 6.00 |
| SCBI | forest | temperate broadleaf or mixed forest | 200 | 0.20 | 7.37 |
| SERC | forest | temperate broadleaf or mixed forest | 200 | 0.22 | 10.25 |
| DSNY | grassland | temperate grassland, savanna and shrubland | 200 | 0.24 | 15.11 |
| JERC | forest | temperate coniferous forest | 200 | 0.19 | 6.01 |
| OSBS | forest | temperate coniferous forest | 200 | 0.24 | 2.66 |
| UNDE | forest | temperate broadleaf or mixed forest | 200 | 0.13 | 13.05 |
| KONZ | grassland | temperate grassland, savanna and shrubland | 200 | 0.53 | 18.84 |
| UKFS | forest | temperate broadleaf or mixed forest | 200 | 0.44 | 38.96 |
| MLBS | forest | temperate broadleaf or mixed forest | 205 | 0.28 | 33.90 |
| ORNL | forest | temperate broadleaf or mixed forest | 180 | 0.36 | 8.08 |
| DELA | forest | temperate broadleaf or mixed forest | 200 | 0.27 | 9.71 |
| DCFS | grassland | temperate grassland, savanna and shrubland | 200 | 0.27 | 26.07 |
| NOGP | grassland | temperate grassland, savanna and shrubland | 200 | 0.47 | 21.44 |
| WOOD | grassland | temperate grassland, savanna and shrubland | 200 | 0.30 | 12.37 |
| STEI | forest | temperate broadleaf or mixed forest | 200 | 0.30 | 10.69 |
| TREE | forest | temperate broadleaf or mixed forest | 200 | 0.78 | 10.45 |
| LENO | forest | temperate broadleaf or mixed forest | 200 | 0.35 | 26.14 |
| TALL | forest | temperate coniferous forest | 200 | 0.22 | 8.65 |
| SRER | shrubland | desert or xeric shrubland | 203 | 0.06 | 4.85 |
| ABBY | forest | temperate coniferous forest | 200 | 0.87 | 28.66 |
| WREF | forest | temperate coniferous forest | 200 | 2.07 | 16.78 |
| SJER | grassland | temperate grassland, savanna and shrubland | 170 | 0.13 | 5.39 |
| SOAP | forest | temperate coniferous forest | 213 | 0.71 | 19.32 |

| | | | | | |
|---|---|---|---|---|---|
| BONA | forest | boreal forest/taiga | 193 | 0.32 | 49.50 |
| DEJU | forest | boreal forest/taiga | 300 | 0.79 | 16.31 |
| HEAL | grassland | tundra | 84 | 2.52 | 45.43 |

**Table S3**. The total stock of SOC in a soil profile including both organic and mineral horizons was related to whole-profile root C (Figure 1; $p < 0.001$, $\chi^2 = 34.46$, adj $r^2 = 0.39$, $n = 43$) and was best predicted by fine-root C (frc), mean annual temperature (MAT), clay (mean% clay) and ecosystem type (land_cov). Of these covariates, land cover was the only non-significant term ($P = 0.55$, $\chi^2 = 2.11$), but was retained in the reduced model based on overall model fit. Whole profile SOC stocks increased with fine-root C stocks ($P = 0.004$, $\chi^2 = 8.26$) and percent clay ($P = 0.001$, $\chi^2 = 10.47$), and declined with MAT ($P = 0.003$, $\chi^2 = 8.57$). A model that lacked root C but included the other covariates had weaker data-model fit than the model with root C ($P = 0.004$, $\chi^2 = 8.49$). Full model: SOC ~ frc+mat+map +clay+land_cov + (1|layer_bot_max). Layer_bot_max = maximum measurement depth. Selected model is highlighted but other models are also shown.

| Model, SOC ~ | AIC | Marginal Pseudo-$R^2$ |
|---|---|---|
| [FULL] frc+mat+map +clay+land_cov + (1\|layer_bot_max) | 836.94 | 0.39 |
| frc+mat+clay+land_cov + (1\|layer_bot_max) | 839.72 | 0.39 |
| mat+map+clay+land_cov + (1\|layer_bot_max) | 845.58 | 0.34 |
| mat+clay+land_cov + (1\|layer_bot_max) | 849.16 | 0.33 |
| frc+map+clay+land_cov + (1\|layer_bot_max) | 853.03 | 0.31 |
| frc+land_cov + (1\|layer_bot_max) | 873.30 | 0.25 |
| frc+mat+clay + (1\|layer_bot_max) | 874.40 | 0.37 |
| frc+clay + (1\|layer_bot_max) | 893.00 | 0.27 |
| frc+mat + (1\|layer_bot_max) | 893.29 | 0.32 |
| frc+map + (1\|layer_bot_max) | 905.19 | 0.25 |

**Table S4a.** Whole profile total root and soil C for the sites with organic horizons only (n = 17): SOC ~ frc+mat+land_cov + map + (1|layer_bot_max). Clay was excluded due to lack of values. Highlighted model is the one reported in the main text (usually selected based on lower AIC; exceptions where less parameters resulted in a similar AIC). AICc was calculated here due to the low number of sites included in the analysis. In the reported best fit model, only root C was significant (p=0.02, $\chi^2$= 5.12), while MAT (p=0.27, $\chi^2$=1.19) and land cover (p<0.92, $\chi^2$= 0.17) were not significant. See caption of Table S3 for model acronyms.

| Model, SOC ~ | AICc | Marginal Pseudo-R2 |
|---|---|---|
| [FULL] frc+mat +land_cov + map (1\|layer_bot_max) | 285.35 | 0.42 |
| frc+mat +land_cov + (1\|layer_bot_max) | 280.47 | 0.42 |
| frc+land_cov + (1\|layer_bot_max) | 279.74 | 0.44 |
| land_cov + (1\|layer_bot_max) | 287.06 | 0.40 |
| frc+ land_cov + (1\|layer_bot_max) | 286.54 | 0.39 |
| frc + mat + (1\|layer_bot_max) | 307.47 | 0.38 |
| frc+ (1\|layer_bot_max) | 315.43 | 0.35 |
| mat + (1\|layer_bot_max) | 312.63 | 0.01 |
| map + (1\|layer_bot_max) | 323.11 | 0.00 |
| land_cover + (1\|layer_bot_max) | 287.06 | 0.40 |

**Table S4b.** Mixed-effects models Full model (mineral horizon only n = 43): SOC ~ frc+mat+map +clay+land_cov + (1|layer_bot_max). Highlighted model is the one reported in the main text (usually selected based on lower AIC; exceptions where less parameters resulted in a similar AIC). Although land cover was included in the best model, it was not a significant predictor of SOC (P = 0.56, $\chi^2$= 2.04), whereas root C (P = 0.04, $\chi^2$= 4.40), MAT (P = 0.04, $\chi^2$= 4.34), and percent clay (P = 0.007, $\chi^2$= 7.36) were significant covariates of SOC. See caption of Table S3 for model acronyms.

| Model, SOC ~ | AIC | Marginal Pseudo-R2 |
|---|---|---|
| [FULL] frc+mat+map +clay+land_cov + (1\|layer_bot_max) | 831.01 | 0.30 |
| frc+mat+clay+land_cov + (1\|layer_bot_max) | 833.71 | 0.30 |
| mat+map+clay+land_cov + (1\|layer_bot_max) | 839.35 | 0.25 |
| mat+clay+land_cov + (1\|layer_bot_max) | 842.74 | 0.24 |
| frc+map+clay+land_cov + (1\|layer_bot_max | 846.22 | 0.22 |
| frc+land_cov + (1\|layer_bot_max) | 863.25 | 0.17 |
| frc+mat+clay + (1\|layer_bot_max) | 868.21 | 0.29 |
| frc+clay + (1\|layer_bot_max) | 885.19 | 0.18 |
| frc+mat + (1\|layer_bot_max) | 883.64 | 0.24 |
| frc+map + (1\|layer_bot_max) | 894.92 | 0.17 |

**Table S5a.** $\beta_{SOC}$ could be predicted using an interaction term between FRC and land cover (P = 0.01, $\chi^2$= 16.61, $r^2$ = 0.32; Figure 3, n = 37). Since only forests and grasslands had enough sites to compare land covers, we reran this analysis removing cultivated lands (one site) and shrublands (three sites) and the interaction between land cover and FRC remained significant (P = 0.02, $\chi^2$ = 10.42, $r^2$ = 0.25, n = 33). Table below shows models explaining $\beta_{SOC}$~ $\beta_{FRC}$ in forests (n = 21) and grasslands (n = 12) for the whole profile (i.e., organic and mineral horizons). Models were evaluated separately for forests and grasslands due to high collinearity between land cover type and $\beta_{FRC}$. Forest models were unable to explain any variation but grassland $\beta_{SOC}$ was significantly related to $\beta_{FRC}$.

| Land cover | Model, beta_SOC ~ | AICc | Marginal Pseudo-R2 |
|---|---|---|---|

| | | | |
|---|---|---|---|
| Forest | [FULL] $\beta_{FRC}$ + mat + map + clay + (1\|layer_bot_max) | -50.21 | 0.07 |
| Forest | $\beta_{FRC}$ + mat + map + (1\|layer_bot_max) | -70.08 | 0.07 |
| Forest | mat + map + clay + (1\|layer_bot_max) | -57.14 | 0.02 |
| Forest | mat + map + (1\|layer_bot_max) | -76.27 | 0.02 |
| Forest | $\beta_{FRC}$ + mat + (1\|layer_bot_max) | -96.43 | 0.07 |
| Forest | $\beta_{FRC}$ + (1\|layer_bot_max) | -113.42 | 0.05 |
| Grassland | [FULL] $\beta_{FRC}$ + mat + map + clay + (1\|layer_bot_max) | 19.33 | 0.36 |
| Grassland | $\beta_{FRC}$ + mat + map + (1\|layer_bot_max) | -6.79 | 0.31 |
| Grassland | mat + map + clay + (1\|layer_bot_max) | 5.47 | 0.14 |
| Grassland | mat + map + (1\|layer_bot_max) | -16.16 | 0.14 |
| Grassland | $\beta_{FRC}$ + mat + (1\|layer_bot_max) | -39.49 | 0.04 |
| Grassland | $\beta_{FRC}$ + (1\|layer_bot_max) | -56.44 | 0.37 |

**Table S5b.** Mixed effects models testing the relationship between $\beta_{SOC}$ and $\beta_{root}$, mean annual temperature (MAT), and mean annual precipitation (MAP) in forest (n = 21) and grassland (n = 12) mineral soil horizons. Models were evaluated separately for each land cover type.

| Land cover | Model, SOC ~ | AICc | Marginal Pseudo-R2 |
|---|---|---|---|
| Forest | [FULL] βFRC + mat + map + clay + (1\|layer_bot_max) | -11.91 | 0.21 |
| Forest | βFRC + mat + map + (1\|layer_bot_max) | -34.23 | 0.21 |

| | | | |
|---|---|---|---|
| Forest | βFRC + map + (1\|layer_bot_max) | -50.78 | 0.12 |
| Forest | βFRC + mat + (1\|layer_bot_max) | -59.77 | 0.22 |
| Forest | βFRC + clay + mat + (1\|layer_bot_max) | -39.11 | 0.23 |
| Forest | βFRC + clay + map + (1\|layer_bot_max) | -30.23 | 0.14 |
| Forest | mat + map + (1\|layer_bot_max) | -41.41 | 0.21 |
| Forest | mat + map + clay + (1\|layer_bot_max) | -20.69 | 0.23 |
| Forest | βFRC + (1\|layer_bot_max) | -74.73 | 0.00 |
| Forest | mat + (1\|layer_bot_max) | -66.39 | 0.23 |
| Forest | map + (1\|layer_bot_max) | -66.39 | 0.23 |
| Forest | clay + (1\|layer_bot_max) | -61.25 | 0.00 |
| Grassland | [FULL] βFRC + mat + map + clay + (1\|layer_bot_max) | 34.25 | 0.22 |
| Grassland | βFRC + mat + map + (1\|layer_bot_max) | 3.31 | 0.21 |
| Grassland | βFRC + mat + (1\|layer_bot_max) | -27.68 | 0.27 |
| Grassland | βFRC + mat + clay + (1\|layer_bot_max) | -4.53 | 0.27 |
| Grassland | βFRC + map + (1\|layer_bot_max) | -19.97 | 0.31 |
| Grassland | βFRC + map + clay + (1\|layer_bot_max) | 3.83 | 0.37 |
| Grassland | mat + map + clay + (1\|layer_bot_max) | 14.41 | 0.15 |
| Grassland | mat + map + | -9.19 | 0.15 |

| | | | |
|---|---|---|---|
| | (1\|layer_bot_max) | | |
| Grassland | βFRC + (1\|layer_bot_max) | -48.61 | 0.31 |
| Grassland | mat + (1\|layer_bot_max) | -33.36 | 0.05 |
| Grassland | map + (1\|layer_bot_max) | -25.33 | 0.05 |

Table S6. Best predictors of inferred priming (residuals from the 1:1 line) across combined ecosystem type, forest or grassland land covers; for both shallow or deep soil layers. We show six models with the lowest AIC out of 40 possible models (of up to 4 predictors).

| Depth | Ecosystem type | Model Parameters | RSquare | RMSE | AICc |
|---|---|---|---|---|---|
| Shallow | all | Ecosystem type, elevation, clay | 0.35 | 0.81 | 80.94 |
| Shallow | all | Ecosystem type, MAT, clay, latitude | 0.40 | 0.79 | 81.50 |
| Shallow | all | Ecosystem type, K, clay | 0.32 | 0.83 | 82.39 |
| Deep | all | Ca, K, pH | 0.42 | 0.47 | 45.92 |
| Deep | all | Ecosystem type, Ca, Mg, K | 0.49 | 0.44 | 46.59 |
| Deep | all | Ecosystem type, Ca, K | 0.48 | 0.44 | 46.74 |
| Shallow | forest | K | 0.14 | 1.26 | 87.47 |
| Shallow | forest | clay | 0.10 | 1.29 | 88.76 |
| Shallow | forest | MAT, latitude | 0.29 | 1.17 | 85.61 |
| Shallow | forest | K, clay | 0.29 | 1.17 | 85.69 |
| Shallow | forest | clay, MAT, latitude | 0.47 | 1.04 | 81.72 |
| Shallow | forest | Na, MAT, latitude | 0.44 | 1.07 | 82.94 |
| Deep | forest | MAT | 0.19 | 1.02 | 77.05 |
| Deep | forest | latitude | 0.15 | 1.05 | 78.41 |
| Deep | forest | aridity index, MAT | 0.24 | 1.02 | 78.53 |
| Deep | forest | Na, MAT | 0.23 | 1.02 | 78.81 |
| Deep | forest | aridity index, MAT, MAP | 0.31 | 0.98 | 78.96 |
| Deep | forest | Mg, Na, MAT | 0.26 | 1.03 | 81.04 |
| Shallow | grassland | Mg | 0.47 | 0.45 | 21.48 |
| Shallow | grassland | clay | 0.33 | 0.50 | 24.14 |
| Shallow | grassland | Mg, MAP | 0.67 | 0.37 | 20.58 |
| Shallow | grassland | Mg, latitude | 0.65 | 0.38 | 21.18 |
| Shallow | grassland | Mg, aridity index, latitude | 0.75 | 0.34 | 23.56 |
| Shallow | grassland | Mg, clay, MAP | 0.73 | 0.35 | 24.23 |
| Deep | grassland | aridity index | 0.23 | 0.29 | 11.30 |
| Deep | grassland | clay | 0.18 | 0.30 | 11.95 |
| Deep | grassland | clay, latitude | 0.36 | 0.28 | 13.67 |
| Deep | grassland | MAT, MAP | 0.34 | 0.28 | 14.11 |
| Deep | grassland | K, clay, latitude | 0.54 | 0.25 | 15.94 |
| Deep | grassland | clay, MAP, latitude | 0.53 | 0.25 | 16.26 |

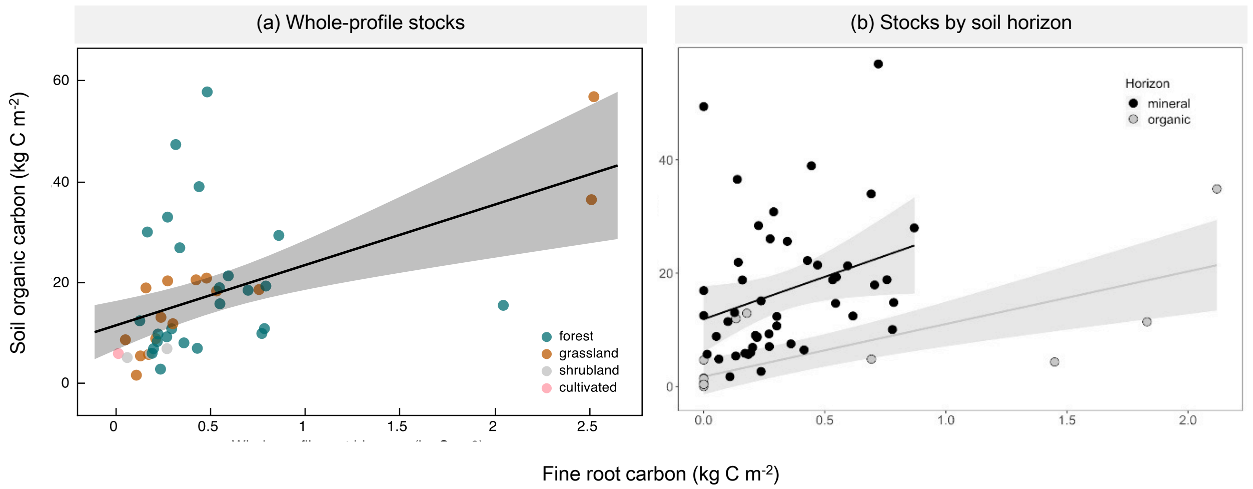


Figure S1. (a) SOC and fine-root biomass C (FRC) stocks, summed across soil horizons, are related across NEON sites (each point represents one NEON megapit at a single site; adj $r^2$= 0.39, $p < 0.001$, n = 43). (b) Horizon-dependent relationships between SOC and fine-root biomass C stocks in organic vs mineral horizons (post-hoc estimated marginal means comparison: p= 0.001, t = 3.85). Points represent megapit samples taken at each NEON site from a given type of horizon (organic or mineral). FRC was the significant predictor of SOC in the organic horizon (adj $r^2$= 0.41, p=0.03, n = 17 out of which 2 were grasslands and rest were forests), while in the mineral horizon other factors such as MAT and percent clay were also important (adj $r^2$= 0.30, p = 0.003, n= 43).

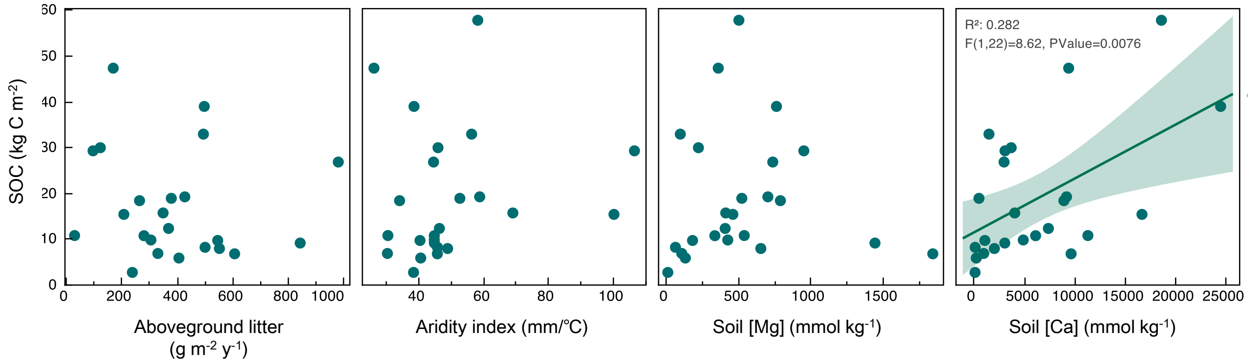


Figure S2. Ecosystem and climate variables exhibiting weak or no relationships with forest SOC. Only soil calcium concentration has a significant linear relationship with SOC.

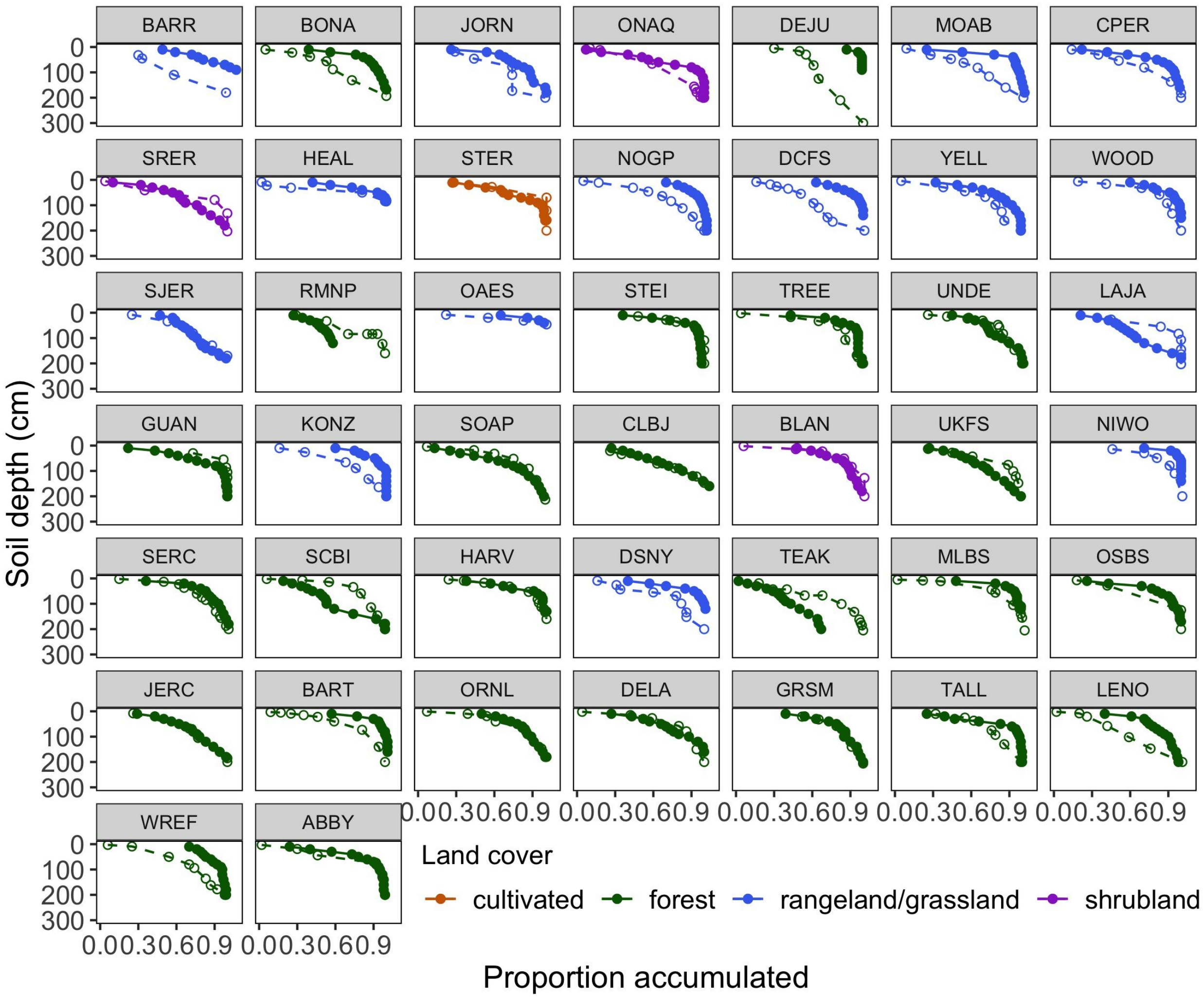


Figure S3. Root and soil carbon accumulation with soil depth at each NEON site. Dashed lines and open circles indicate SOC proportion accumulated and solid lines and filled circles indicate root biomass C. Facets are arranged by order of increasing MAP beginning with BARR having the lowest MAP (105 mm/y) to ABBY with the highest MAP (2451 mm $y^{-1}$).

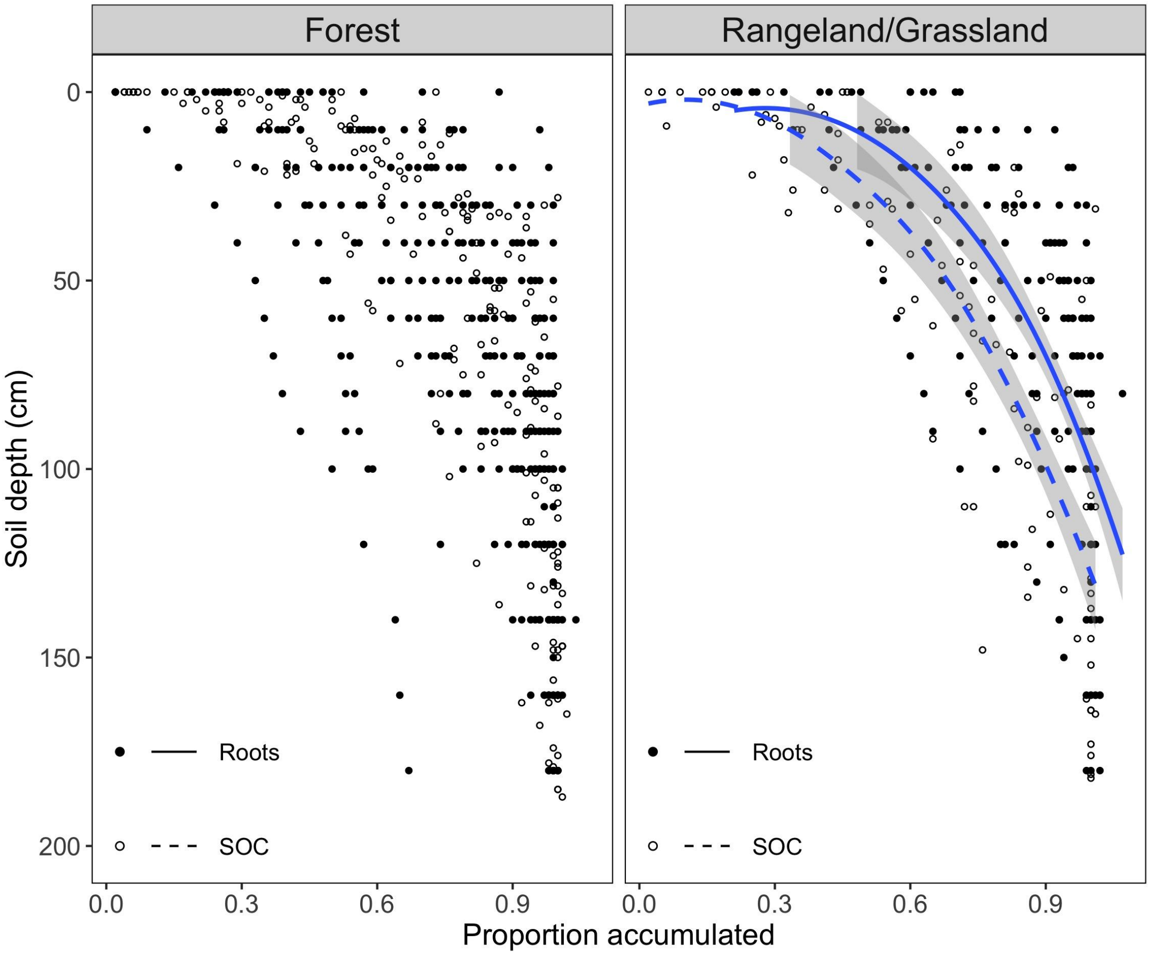


Figure S4. Average cumulative proportion of FRC and SOC across depth intervals in forests (n= 25 and excludes 4 sites where beta SOC could not be calculated due to too few measurements in the profile as detailed in methods) and grasslands (n = 12). Points represent the proportion of either FRC (filled circles) or SOC (open circles) accumulated within a depth interval at each site (See Figure S3 for site-specific accumulation profiles). Depth intervals are plotted with respect to the top of the depth interval (e.g., the interval 0 - 10 cm is plotted at 0 cm). Lines represent the smoothed, statistically significant regression and gray shaded areas represent the 95% confidence interval. The average beta coefficient for root biomass C in forests was 0.958 and in grasslands was 0.942, and the average beta coefficient for SOC in forests was 0.974 and in grasslands was 0.978. Using these root beta values, we also calculated the depth at which 50% or 95% of the root biomass occur. For forests, 50% of the roots occurred above 91 cm and 95% of the roots occurred above 106 cm. Conversely, for grasslands, 50% of the roots occurred above 65 cm and 95% occurred above 76 cm.

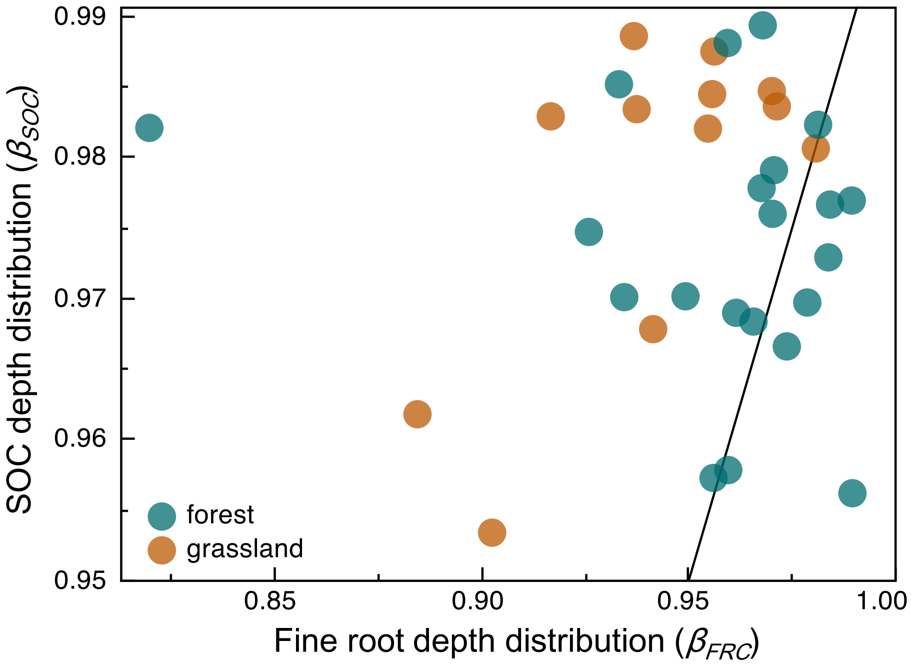


Figure S5: Fine-root biomass C and SOC depth profiles (inferred from β coefficients) are unrelated across land cover types.The black line represents a 1:1 relationship between $\beta_{FRC}$ and $\beta_{SOC}$. Points represent NEON megapit samples and are colored by land cover. Within land covers, forests show high variability while grassland $\beta_{SOC}$ can be predicted by $\beta_{FRC}$ (Table S5a and b). For whole profiles (organic+mineral), $\beta$ coefficients describing the depth profiles of SOC and FRC were unrelated despite inclusion of MAT, MAP or clay in predictive models. However, SOC $\beta$ could be predicted using an interaction term between FRC and land cover ($P = 0.01$, $\chi^2 = 16.61$, $r^2 = 0.32$; Figure 3, $n = 37$). Since only forests and rangeland/grasslands had enough sites to compare land covers, we compared each land cover's predictive models (Table S5a) and found that forest $\beta_{SOC}$ had high variability and low model predictability. Conversely, grassland $\beta_{SOC}$ had lower variability and could be predicted using root $\beta$ (Table S5a). Next, we explored whether high root biomass in the organic layer may be biasing the $\beta_{FRC}$ to be much more shallowly distributed than $\beta_{SOC}$. Thus, a stronger root-soil beta relationship could exist if we examined only the mineral horizons. Indeed we found stronger predictive models of mineral horizons but forest variability remained high (Table S5b). We found that similar to the whole-profile results, $\beta_{SOC}$ from only mineral horizons could be explained with a function that included $\beta_{FRC} \times$ land cover but unlike the whole-profile, this model also included %clay ($p = 0.03$, $X^2 = 10.67$, $R^2 = 0.28$, $n = 33$). Here again, the correlation in mineral soils between $\beta_{SOC}$ and $\beta_{FRC}$ coefficients was higher in rangelands/grasslands than in forests ($p = 0.02$, $t = -2.44$). Climate variables MAT and MAP did not co-vary with $\beta_{SOC}$ ($p > 0.05$). Thus, overall we find that, contrary to our hypothesis, the depth profiles (inferred using exponential decay functions and resulting beta coefficient; $\beta$) of FRC and SOC were not related but this relationship improved if organic horizon data was removed from the analysis. Our analyses again highlighted differences among forests and grasslands, particularly high variability in forest SOC $\beta$ where climatic and soil factors were more important predictors than root $\beta$. Conversely, in grasslands, root and SOC $\beta$ were related. We found no significant predictors of forest $\beta_{SOC}$ distributions, except aboveground inputs (Figure S6).

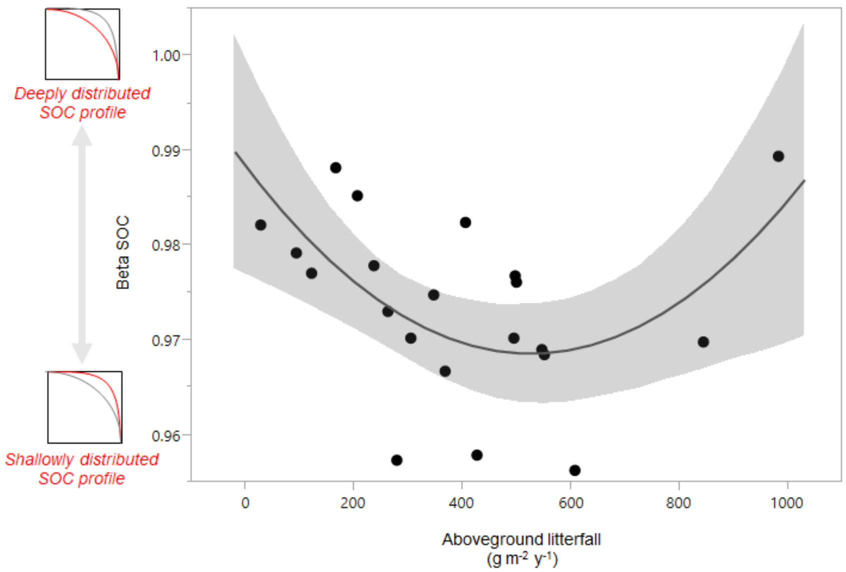


Figure S6. SOC depth distributions($\beta_{SOC}$) at NEON forest sites are significantly related to aboveground litter fall (Quadratic regression: $\beta_{SOC}$ = 0.9 - 1.9e-5 * litter + 7.2e-8 * (litter-394.3)$^2$; $r^2$=0.33, p=0.02). Without the two largest aboveground litter values, the regression is still significant ($r^2$= 0.30, p = 0.014) and supports a linear negative relationship between litter and $\beta_{SOC}$.

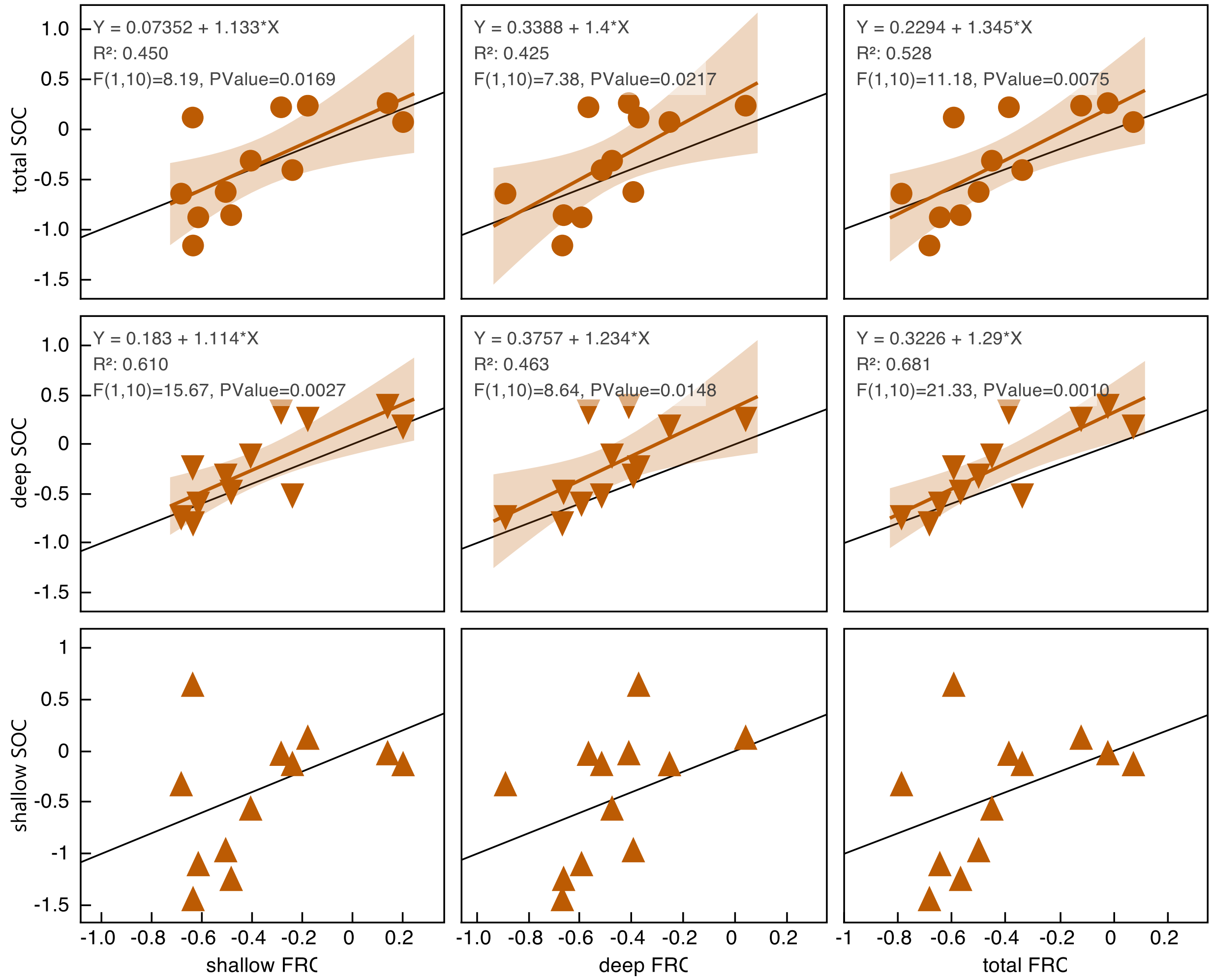


Figure S7: Relationships between shallow, deep and total FRC and shallow, deep and total SOC (shallow = 0 to 30 cm, deep = 30 to profile bottom, total = whole profile). Only grassland data points are shown (3 tundra sites removed). Note that data have been standardized ([raw data value-mean value]/standard deviation] and therefore are unitless. Black line is the 1:1 line, such that values under the line denote increased FRC decreasing SOC (priming effect) and values above the line denote increased FRC increasing SOC (net SOC accrual). Linear regression results are shown when significant ($p<0.05$).

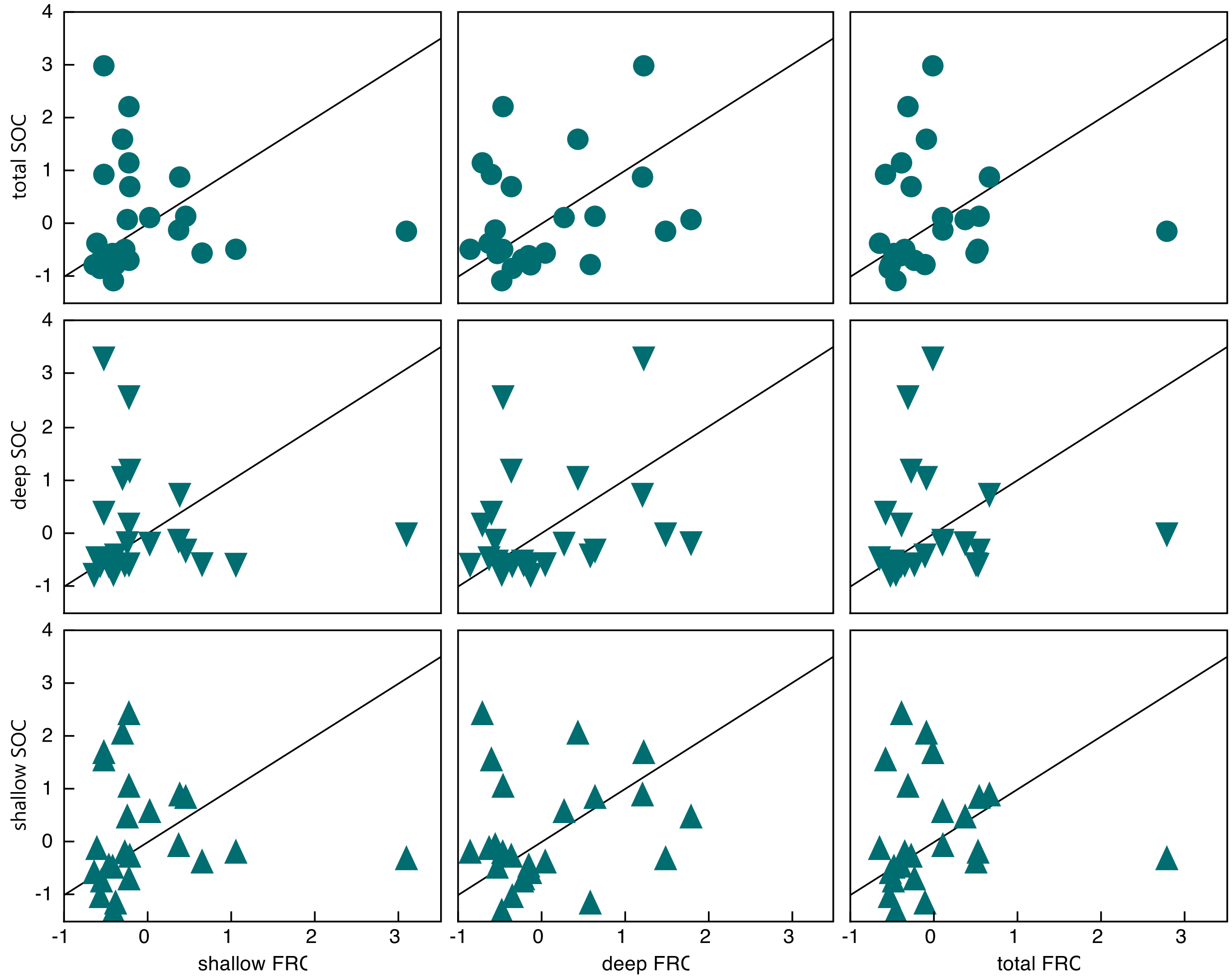


Figure S8. Relationships between shallow, deep and total FRC and shallow, deep and total SOC (shallow = 0 to 30 cm, deep = 30 to profile bottom, total = whole profile). Only forest data points are shown. Note that data have been standardized ([raw data value-mean value]/standard deviation] and therefore are unitless. Black line is the 1:1 line. Values under the line denote increased FRC decreasing SOC (inferred priming effect) and values above the line denote increased FRC increasing SOC (inferred net accrual). No significant relationships ($P>0.05$) are seen even if the outlier with the highest root values (WREF site) is removed.

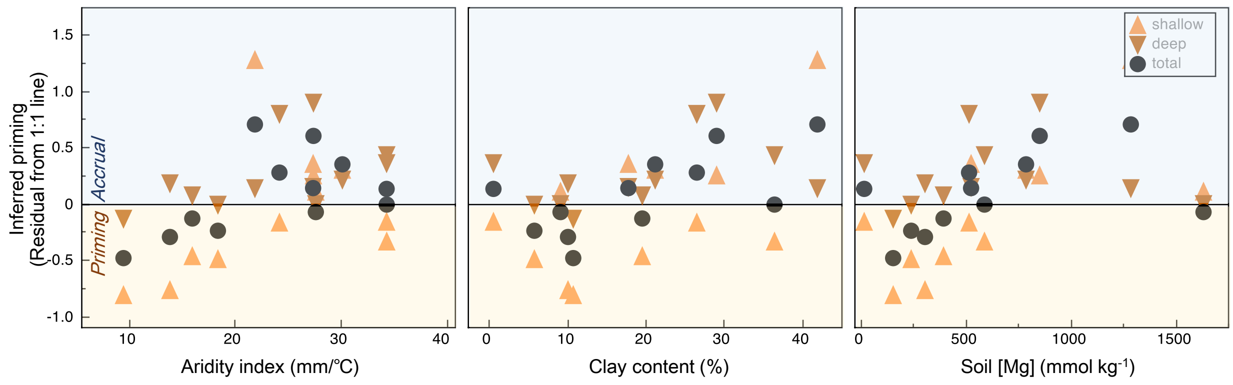


Figure S9. Best predictors of grassland inferred priming. Aridity, clay and micronutrients drive priming (inferred using the residuals of the 1:1 fit between FRC and SOC), although only certain soil depths are significant for each driver in linear regressions. Aridity across soil layers (total) is related to inferred priming ($r^2$= 0.29, p=0.07). Shallow ($r^2$= 0.33, p=0.05) and total ($r^2$= 0.43, p=0.02) clay content are related to inferred priming. Lastly, soil magnesium concentration in shallow layers is related to inferred priming ($r^2$= 0.47, p=0.01). Values above zero (blue region) corresponds to inferred net SOC accrual, and values below zero (yellow region) corresponds to net inferred priming.

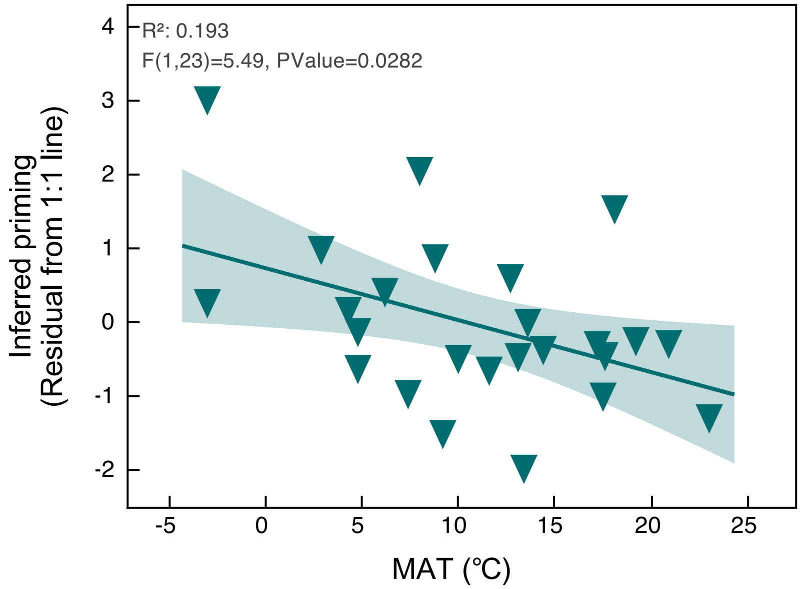


Figure S10. Warmer forests have more inferred priming in deep soil layers.

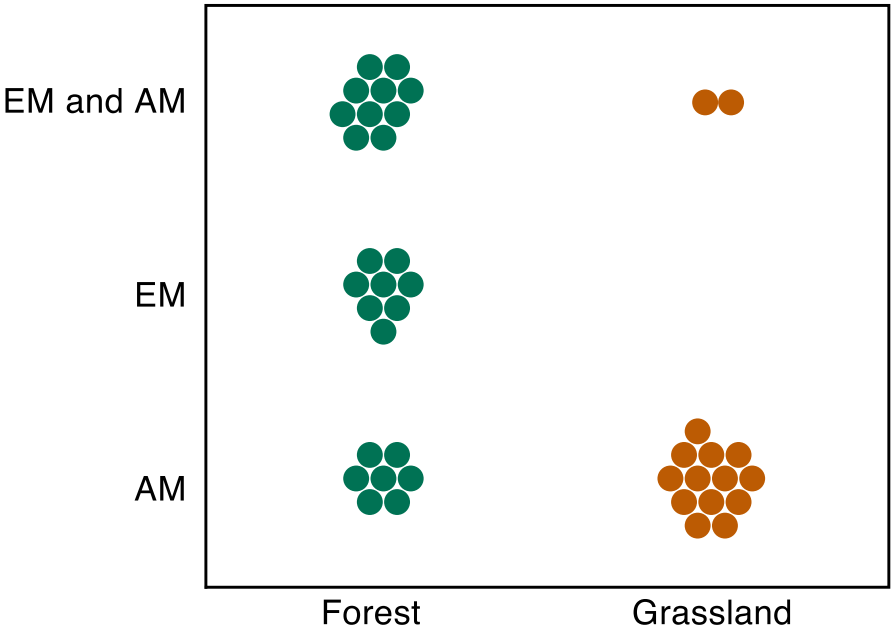


Figure S11: Dominant mycorrhizal types at each of the forest and grassland sites. Two dominant types of mycorrhizal fungi, ectomycorrhizal (EM) fungi and arbuscular mycorrhizal (AM) fungi were determined based on dominant species reported in NEON [54] that were used to infer mycorrhizal types based on [48]

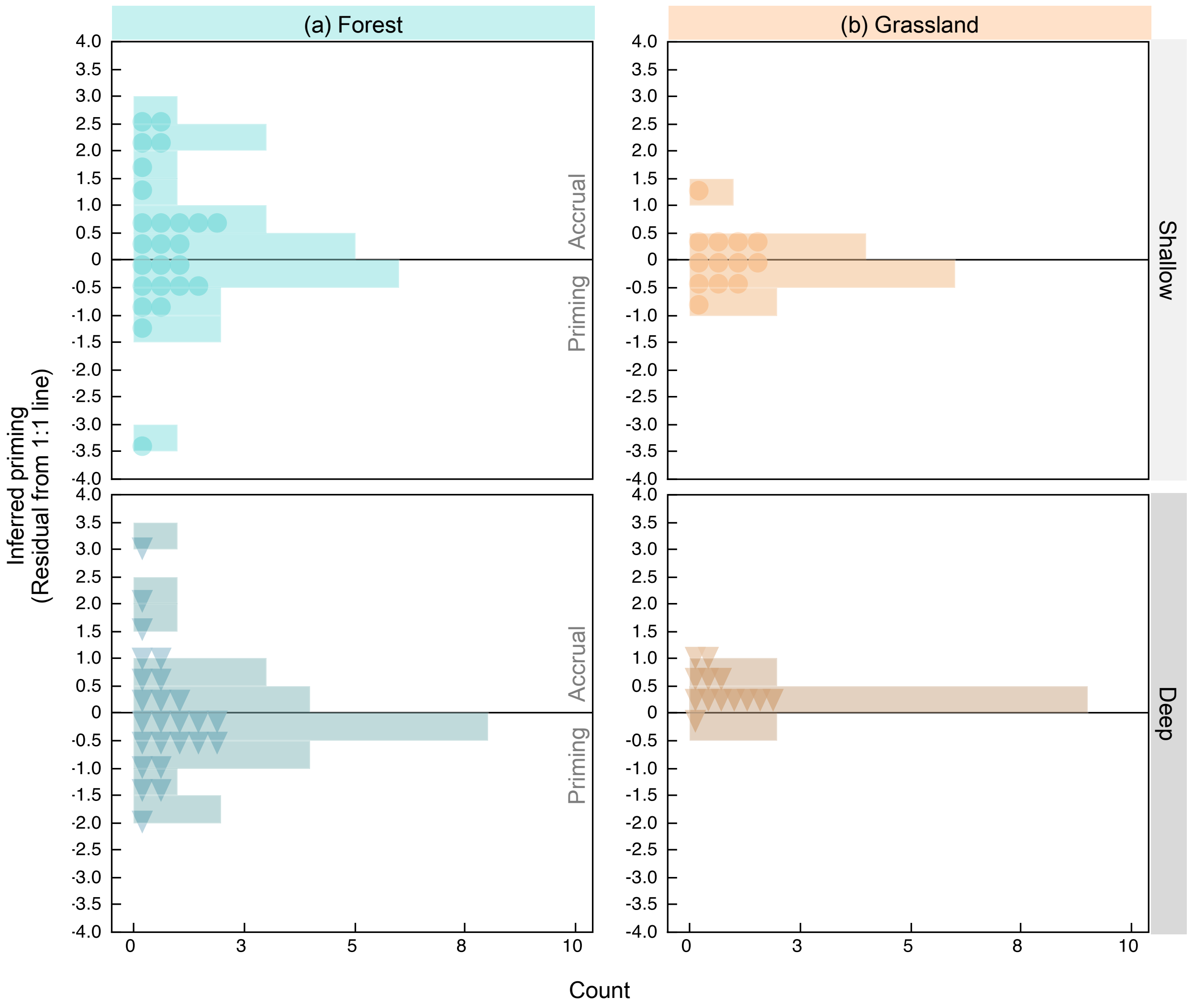


Figure S12. Raw data points (corresponding to Figure 3) and distributions of inferred priming (residuals from the 1:1 line between standardized FRC and SOC; see Figure 2). Data are shown for shallow (<30 cm depth) and deep (30-200 cm) layers from both (a) forest and (b) grassland sites. Bins above the zero line are reflective of sites with possible SOC accrual and bins below the zero line are priming.
